# The Hunchback temporal transcription factor determines interneuron molecular identity, morphology, and presynapse targeting in the *Drosophila* NB5-2 lineage

**DOI:** 10.1101/2024.10.07.616945

**Authors:** Heather Q. Pollington, Chris Q. Doe

## Abstract

Interneuron diversity within the central nervous system (CNS) is essential for proper circuit assembly. Functional interneurons must integrate multiple features, including combinatorial transcription factor (TF) expression, axon/dendrite morphology, and connectivity to properly specify interneuronal identity. Yet, how these different interneuron properties are coordinately regulated remains unclear. Here we used the *Drosophila* neural progenitor, NB5-2, known to generate late-born interneurons in a proprioceptive circuit, to determine if the early-born temporal transcription factor (TTF), Hunchback (Hb), specifies early-born interneuron identity, including <u>molecular profile</u>, <u>axon/dendrite morphology</u>, and <u>presynapse targeting</u>. We found that prolonged Hb expression in NB5-2 increases the number of neurons expressing early-born TFs (Nervy, Nkx6, and Dbx) at the expense of late-born TFs (Runt and Zfh2); thus, Hb is sufficient to promote interneuron molecular identity. Hb is also sufficient to transform late-born neuronal morphology to early-born neuronal morphology. Furthermore, prolonged Hb promotes the relocation of late-born neuronal presynapses to early-born neuronal presynapse neuropil locations, consistent with a change in interneuron connectivity. Finally, we found that prolonged Hb expression led to defects in proprioceptive behavior, consistent with a failure to properly specify late-born interneurons in the proprioceptive circuit. We conclude that the Hb TTF is sufficient to specify multiple aspects of early-born interneuron identity, as well as disrupt late-born proprioceptive neuron function.

## Introduction

During development, the generation of neuronal diversity is a crucial component in assembling proper neural circuit assembly and increases the complexity of neural circuitry and behavioral output. The central nervous system (CNS) consists of a diverse array of neurons that differ in transcription factor and neurotransmitter expression, morphology, connectivity, and electrophysiology. All these features are integrated to define unique neuron identities, enabling each neuron to form specific and stereotyped neural circuits that generate appropriate behaviors.

Neurogenesis in both Drosophila and vertebrates is initiated by spatial cues that specify progenitor identity [1], followed by temporally expressed transcription factors (TTFs) that diversify the progeny of each progenitor [2–4]. In the mouse retina, retinal progenitor cells (RPCs) sequentially generate multiple cell types necessary for retinal circuit assembly [5]. RPCs express the TTF cascade: Ikaros > Pou2f1/Pou2f2 > FoxN4 > Casz1, each specifying specific cell types [4]. Ikaros expression is maintained in early-born RPC progeny and promotes expression of the homeodomain transcription factor, Prox-1, required to specify early-born horizonal cell fate [6]. In the developing mammalian cortex, apical radial glia (aRGs) produce cortical neurons that settle in functionally distinct laminar layers in an inside-out fashion, generating deep layered early-born neurons and superficially located late-born neurons [3,4]. As in the retina, Ikaros is expressed early in neuronal progenitors of the mouse neocortex [7]. Prolonged Ikaros expression in cortical progenitors results in an increase in the number of early-born neurons expressing the early-born cortical TFs, Tbr1 and Foxp2, where it is required for early-born neuronal fate [7]. In the zebrafish spinal cord, early-born motor neurons and interneurons form a fast-swimming circuit, while later-born neurons form a slow-swimming circuit [8]. Early-born spinal cord neurons also show distinct morphological and connectivity differences compared to later-born neurons [8]. Thus, temporal patterning within progenitor lineages is an essential mechanism for generating neuronal diversity in the vertebrate CNS.

Similarly, *Drosophila* neural progenitors, called neuroblasts (NBs), sequentially produce motor neurons, interneurons, and glial cell type progeny that settle in a deep-to-superficial pattern [9–12], similar to mammalian cortical development [1]. *Drosophila* NBs generate a stereotyped sequence of neurons and glia in all parts of the CNS, but here we focus on the ventral nerve cord, analogous to the vertebrate spinal cord. NBs in the VNC sequentially express the following TTF cascade: Hunchback (Hb, ortholog of Ikaros) > Krüppel (Kr) > Nubbin/Pdm2 (Pdm) > Castor (Cas; ortholog of Casz1) > Grainy head (Grh) [2]. During each expression window, the NB asymmetrically divides to generate a smaller ganglion mother cell (GMC), which then divides to produce two post-mitotic neurons that transiently maintain the TTF expressed at the time of birth. This generates early-born neurons that express Hb, required for establishing motor neuron identity [9,13–17].

The role of Hb in specifying *Drosophila* motor neuron fate has been most extensively investigated in the NB3-1 and NB7-1 lineages [9,13–17]. NB7-1 generates the U1-U5 motor neurons in sequential birth-order: U1/U2 express Hb, U3 expresses Kr, U4 expresses Pdm, and U5 expresses Pdm/Cas [9]. Prolonged Hb expression in NB7-1 generates increased numbers of U1/U2 at the expense of later-born motor neurons [9,14,15]. The ectopic U1/U2 motor neurons lack expression of the late-born transcription factors, Zfh2, display U1/U2 morphology, and form synapses with U1/U2 dorsal muscle targets [14,15]. Similarly, prolonged Hb expression in NB3-1 increases functional synapses innervating early-born muscle targets and decreases synapse formation to later-born ventral muscle targets [16]. Increased synaptic connections to early-born muscles increases presynaptic vesicle release; however, there is no difference in miniature-excitatory post-synaptic potential (mEPSP) amplitude, suggesting ectopic Hb motor neurons may undergo homeostatic compensation [16].

The role of Hb in generating motor neuron identity and connectivity is well characterized, yet relatively little is known about the role of TTFs in generating <u>interneuron</u> identity. Interneuron and motor neuron axons have different constraints on circuit formation. While motor neuron axons select a muscle target from a small pool of muscles, interneuronal axons must bypass a vast array of potential neuron partners within the synaptically dense neuropil, a much more complex environment, to form appropriate synaptic connections to partner neurons. In addition, interneuron neurites are directed by gradients of the attractant and repulsive extrinsic signaling cues: Semaphorin 1a, 2a, and 2b, Slit, and Netrin [18–22]. Interneurons expressing specific guidance receptors are either attracted or repelled from discrete domains within the neuropil, directing neurites to extend to distinct neuropil regions. These extrinsic cues may contribute to interneuron morphology and connectivity; however, the role of neuron intrinsic mechanisms is not well understood.

To understand whether the mechanisms that Hb uses to specify early-born motor neuron identity is conserved in interneurons, we investigated the role of Hb in specifying interneuron identity in the NB5-2 lineage. NB5-2 predominately generates interneuron progeny, with the late-born interneurons forming a proprioceptive circuit [11,23]. We generated a NB5-2 split Gal4 line to genetically label NB5-2 and its progeny. We found that NB5-2 expresses the canonical TTF cascade and TTF expression is transiently maintained in the neuronal progeny. When we prolonged expression of Hb in the NB5-2 lineage, it resulted in an increased number of Hb+ NB5-2 interneurons that express early-born TFs (Nervy, Dbx, and Nkx6) at the expense of neurons expressing late-born TFs (Runt and Zfh2). In addition, we identified several Hb+ neurons with an axon/dendrite morphology distinct from that of later-born interneurons, forming a unique diagonal contralateral projection. At the synaptic level, ectopic Hb expression in late-born neurons resulted in an increase in synapse number at neuropil locations normally targeted by NB5-2 Hb+ early-born neuronal presynapses. Finally, we found that prolonged NB5-2 Hb expression generates behavioral defects similar to disrupting the late-born proprioceptive circuit: decreased locomotor velocity and increased number of C-shaped body bends. In summary, we show that Hb specifies early-born NB5-2 interneuron molecular identity, axon/dendrite morphology, and presynapse targeting (essential for proper connectivity), and that proper TTF identity is necessary for appropriate behavioral output.

## Results

### Generation of a split-Gal4 line expressed in NB5-2 and its embryonic lineage

To create a transgene specifically expressed in the NB5-2 lineage, we constructed a split-Gal4 based on the intersection of *wingless* (*wg*), expressed in row 5 neuroectoderm, and *ventral nervous system defective* (*vnd*), expressed in medial column neuroectoderm (Figure 1A,B). The intersection of these two hemi-drivers defines the neuroectoderm domain that generates NB5-2, and the split-Gal4 is strongly and specifically expressed in NB5-2 and its progeny (Figure 1C). We refer to this split-Gal4 line alone as NB5-2-Gal4, or NB5-2>GFP when the NB5-2-Gal4 is driving expression of UAS-GFP. We note that NB5-2-Gal4 has occasional expression in the NB5-1 lineage, but this has no effect on our experiments as NB5-1 does not express Hb, Kr, or Pdm and has a small, simple Cas+ lineage with a distinct medial fascicle into the neuropil [24,25]. Thus, we conclude that NB5-2-Gal4 is an appropriate tool for driving gene expression in NB5-2 and its embryonic progeny.

**Figure 1.**
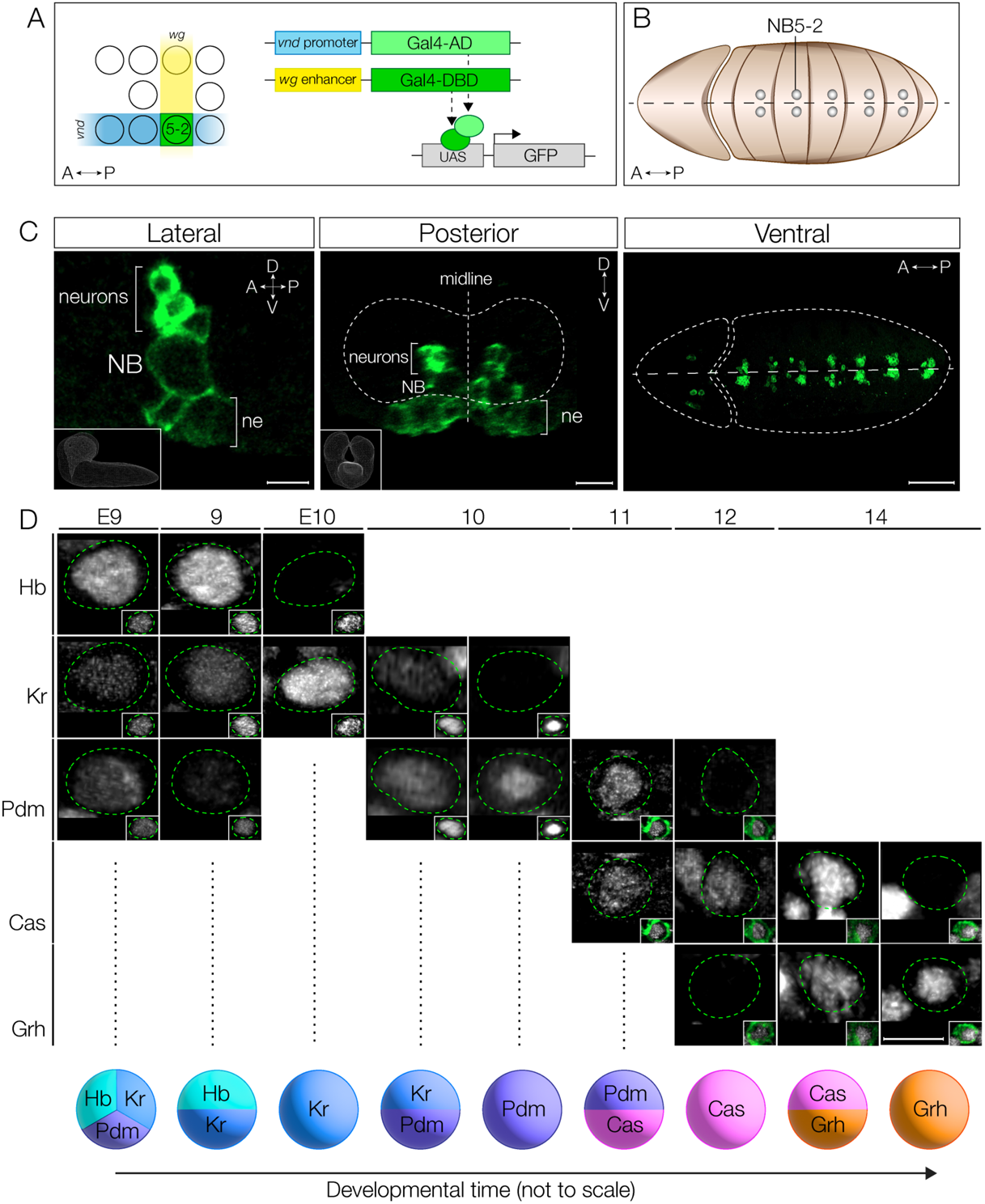
NB5-2 expresses the canonical TTF cascade. (A) Schematic of NB5-2 split Gal4 (green) construction using the unique NB5-2 expression combination of *wingless* (*wg*; yellow) and *ventral nervous system defective* (*vnd*; blue). (B) Schematic of segmentally repeating NB5-2 location in early embryo. Anterior left, ventral view. (C) NB5-2>GFP expression in stage 11 embryo in the lateral (left panel; inset shows CNS volume from lateral view; Scale bar: 5 µm.), posterior (middle panel; inset shows CNS volume from posterior view; Scale bar: 10 µm), and ventral (right panel; Scale bar: 50 µm) views labels NB5-2, NB5-2 neuron progeny, and a small population of neuroepithelium (ne). (D) NB5-2 TTF expression cascade at stages 9-14. NB5-2 identification was achieved using the NB markers Deadpan (Dpn; insets in all expect stg 10) or Worniu (Wor; inset stg 10 only). Due to delayed GFP (green) expression in stage 9 and early stage 10 embryos, NB5-2 was identified as the most medially located NB, anteriorly adjacent to the *engrailed* expression domain (see Supp. Figure 1). A GFP labeled NB5-2 was used to identify NB5-2 after early stage 10. Scale bar: 5 µm.

### NB5-2 expresses the canonical TTF cascade

Most VNC NB lineages sequentially expresses Hb, Kr, Pdm, Cas, and Grh and include intervals where adjacent TTFs are co-expressed [2]. Here we determine whether NB5-2 also expresses this TTF cascade, and whether there are gene expression overlaps within the series. We found that NB5-2>GFP expression was not detected until late stage 10, and thus at earlier stages NB5-2 was identified by its position in the NB array: the most medial row 5 NB just anterior to the row 6/7 Engrailed expression domain (Supp. Figure 1). Using these markers prior to stage 10, and NB5-2>GFP expression beginning at stage 10, we were able to characterize the TTF cascade in NB5-2 throughout embryogenesis (Figure 1D). We found that NB5-2 initially expresses Hb, low levels of Kr, and transient Pdm; brief expression of Pdm during the NB first division window has been previously observed in the NB4-2 lineage and thought to be protein inherited from the Pdm+ neuroectoderm [26]. Subsequently, NB5-2 sequentially expresses Kr, Pdm, Cas, and Grh with a period of overlap in each case (Figure 1D; Supp. Figure 1). We conclude that NB5-2 undergoes the canonical TTF cascade, with transient overlap of each TTF, resulting in nine different TTF expression windows. Most importantly for this work, Hb is detected in the early portion of the NB5-2 lineage.

### NB5-2 neuronal progeny express the TTF present at their time of birth

In most NB lineages, TTF expression in the NB is transiently maintained in the progeny born during each NB TTF expression window [9,12]. Indeed, we find that each TTF can be detected within a subset of NB progeny (Figure 2A-C; quantified in D). Each window of NB expression, both single and dual gene overlap, contributes to the neuronal progeny (Figure 2A-C; quantified in D). Transient expression varied among TTFs: Hb, Kr, and Grh expression was maintained until the L1 larval stage (and possibly longer); Pdm expression was detectable only until embryonic stage 14; and Cas was detectable until late embryonic stage 17, with little expression observed in newly hatched larvae (data not shown). Neurons expressing early TTFs are located in a deep layer closer to the neuropil, while neurons expressing later TTFs are located more superficially (Figure 2A-C), as expected from previous work [9,11,12]. In addition, neurons from sequential TTF windows invariably have adjacent cell body positions (Figure 2A-C).

**Figure 2.**
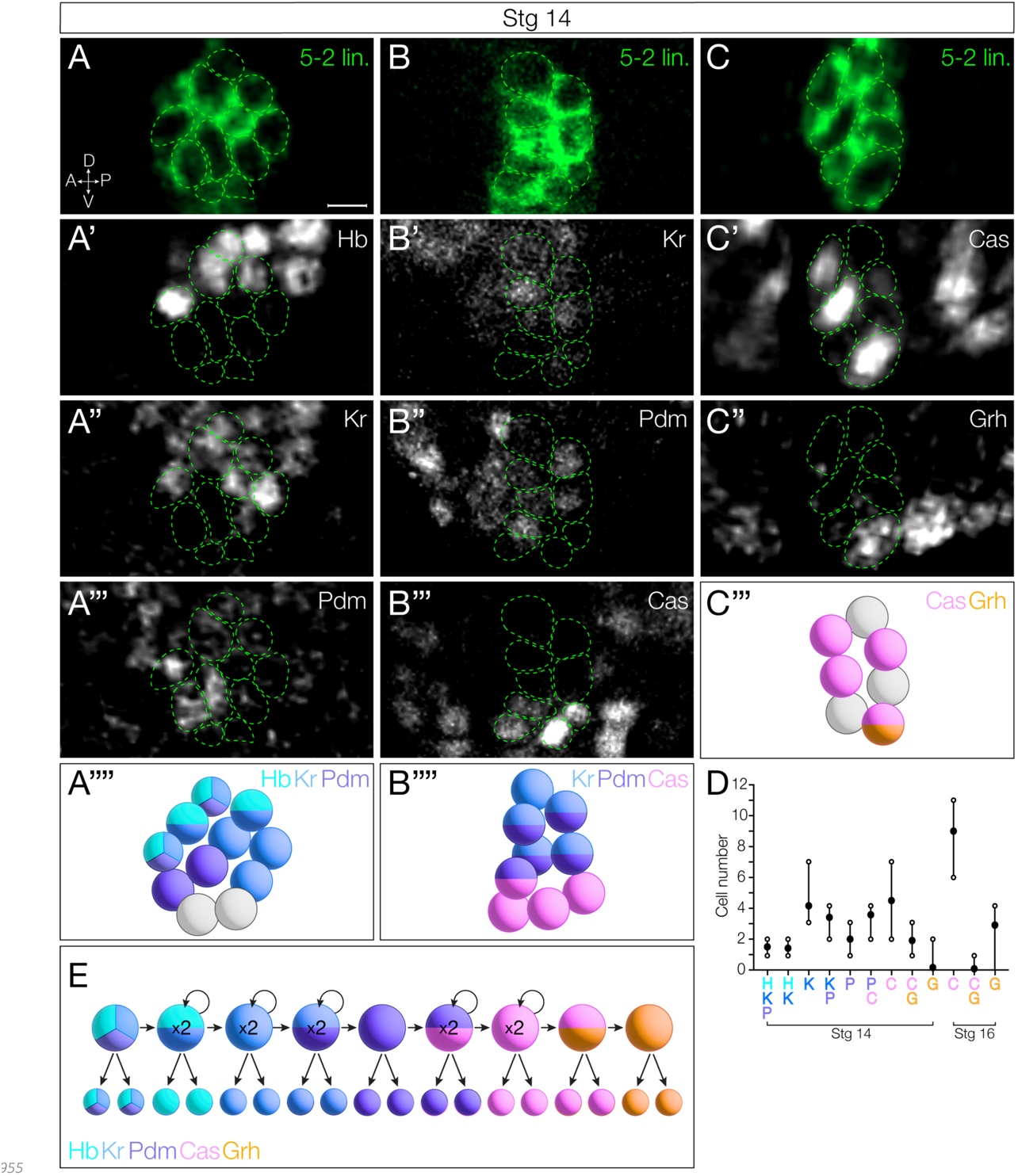
NB5-2 progeny maintain TTF expression present at time of birth. (A) NB5-2>GFP (green) progeny expressing Hb (A’), Kr (A’’), and Pdm (A’’’) at stage 14; A schematic summary of Hb (cyan), Kr (blue), Pdm (purple) expression combinations (A’’’’). Anterior left, lateral view. Scale bar: 4 µm. (B) NB5-2>GFP progeny expressing Kr (B’), Pdm (B’’), and Cas (B’’’); A schematic summary of Kr, Pdm, and Cas (magenta) expression combinations (B’’’’). (C) NB5-2>GFP progeny expressing Cas (C’) and Grh (C’’); A schematic summary of Cas and Grh (orange) expression combinations (C’’’). (D) Quantification of NB5-2 neurons expressing each TTF singly or in combination in stage 14 and 16 embryos. (E) Schematic of NB5-2 divisions during each TTF expression window.

Interestingly, there are different numbers of neurons expressing each TTF (or TTF combination) (Figure 2D), which allows us to use neuron numbers as a proxy for the length of each TTF expression window. For example, at stage 14, Hb is detected in 5 neurons, which suggests that the NB Hb expression persisted for 3 divisions (each division making a Hb+ GMC that makes a pair of sibling neurons). In this way, we infer that Kr alone has a one division window to contribute 2 neurons, Kr/Pdm has one division window (2 neurons), Pdm alone has a one division window (2 neurons), Pdm/Cas has a two division window (4 neurons), Cas alone has a two division window (4 neurons), Cas/Grh has a one division window (2 neurons), and Grh alone has a one division window (2 neurons) (summarized in Figure 2E). Note that the number of Cas+Grh-neurons increases between stage 14 to stage 16 (Figure 2D), consistent with a second window of Cas expression.

### Neuronal settling position is correlated with birth-order

Previous work has shown that early-born Hb+ neurons reside closest to the neuropil, being physically displaced into more dorsal regions by later-born neurons. Conversely, later-born Cas+ neurons take more superficial positions [9–12]. To determine if NB5-2 progeny settling position is correlated with birth-order, we mapped the deep-to-superficial position of the NB5-2 progeny. We found that Hb+ and Kr+ neurons are located in overlapping deep layers (Figure 3A). Pdm neurons are positioned more superficially (Figure 3A-B), with some overlap with both Hb and Kr, consistent with overlapping expression patterns seen in NB5-2 (Figure 1). Lastly, both Cas+ and Grh+ neuron populations are located in the most superficial layers (Figure 3B-C). We conclude that (1) neuronal settling position is correlated with birth-order in the NB5-2 lineage (Figure 3D), and (2) that TTF expression, neuronal settling position, and birth-order are tightly correlated. These findings all support a model in which each TTF is inherited by the neuronal progeny born during each TTF expression window, with little migration or movement, resulting in early-born neurons located in deep layers and late-born neurons located in superficial layers.

**Figure 3.**
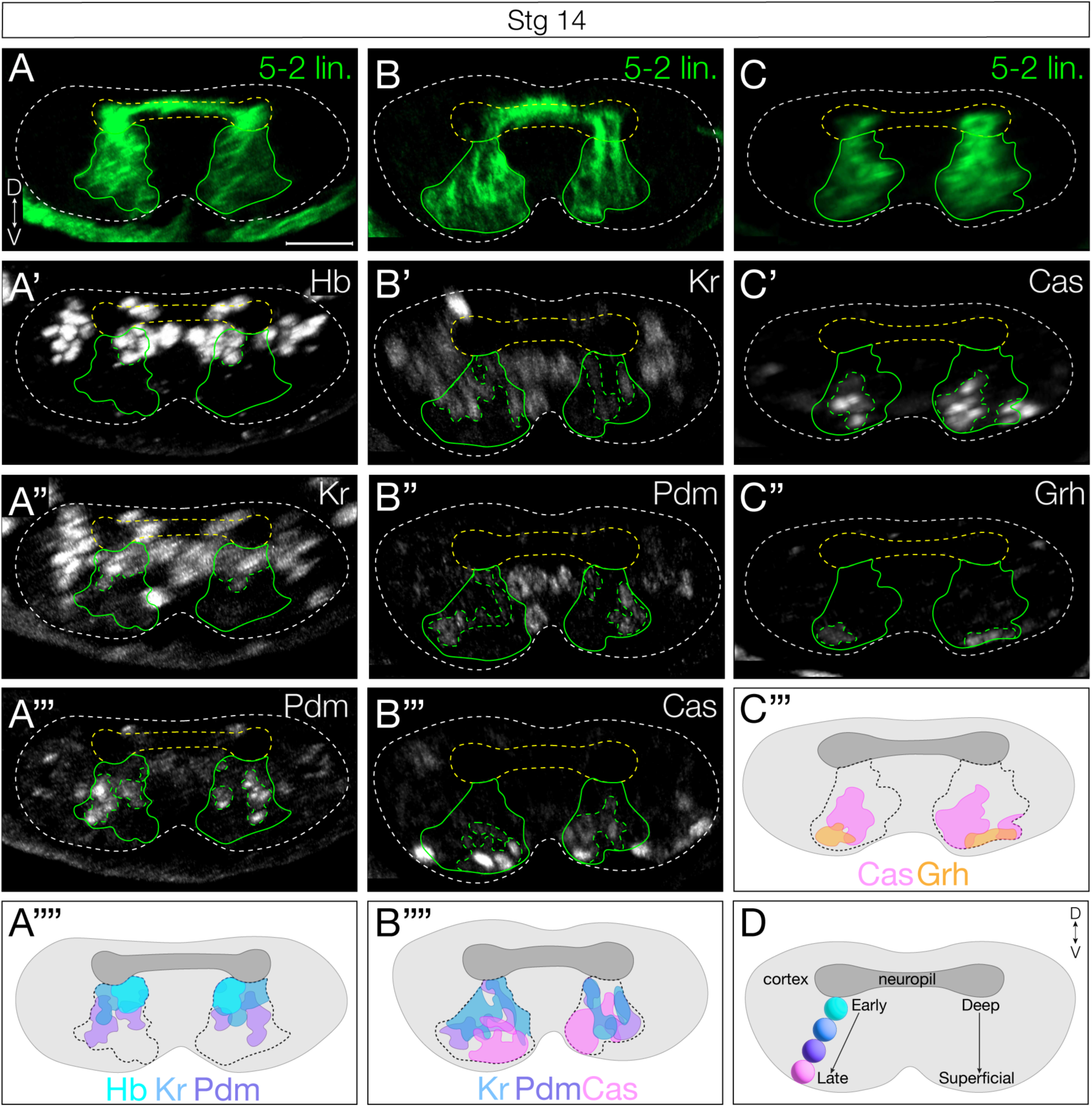
NB5-2 progeny deep-superficial settling position is correlated with birth-order. (A) NB5-2>GFP (green) progeny settling position of neurons expressing Hb (A’), Kr (A’’), and Pdm (A’’’); A schematic summary of Hb (cyan), Kr (blue), and Pdm (purple) settling position overlap (A’’’’). Dorsal up, posterior view. Scale bar: 10 µm. (B) NB5-2>GFP progeny settling position of neurons expressing Kr (B’), Pdm (B’’), and Cas (B’’’); A schematic summary of Kr, Pdm, and Cas (magenta) settling position overlap (B’’’’). (C) NB5-2>GFP progeny settling position of neurons expressing Cas (C’) and Grh (C’’); A schematic summary of Cas and Grh (orange) settling position overlap (C’’’). (D) Cartoon of stereotypical early-to-late born cell body positions within the VNC cortex.

### Prolonged Hunchback expression in NB5-2 prevents late TTF expression

To distinguish the relative importance of birth-order and TTF expression in specifying interneuronal identity, we broke their correlation by mis-expressing Hb in the NB5-2 lineage, which allowed us to determine which -- birth-order or TTF identity -- is more important for neuronal identity. We used our NB5-2-Gal4 line to specifically misexpress Hb throughout the lineage, and assay for alterations in the expression of later-born TTFs. If time of birth is used to generate interneuron diversity, we would expect little effect of Hb misexpression; in contrast, if the TTF identity is causal for interneuron identity, we would expect to see a clear transformation of late-born to early-born interneuron identity. In the following sections we determine the effect of Hb misexpression on interneuronal <u>molecular</u> identity, <u>axon/dendrite morphology</u>, and <u>presynapse targeting</u> within the dense neuropil.

We first validated the ability of Hb misexpression to repress later NB TTFs; in other NB lineages, misexpression of Hb can result in delayed or absent expression of late TTFs [9]. The timing of Hb misexpression is also important: NBs have a relatively short competence window for responding to Hb [27,28]. We found that NB5-2-Gal4 drives misexpression of Hb beginning at early stage 10, soon after the termination of endogenous Hb expression (Figure 4A), and prior to the termination of the competence window at stage 12 [27]. Importantly, we found that misexpression of Hb in the NB5-2 lineage resulted in a highly penetrant loss of expression of all later-born TTFs (Figure 4A; summarized in B), a prerequisite for subsequent experiments.

**Figure 4.**
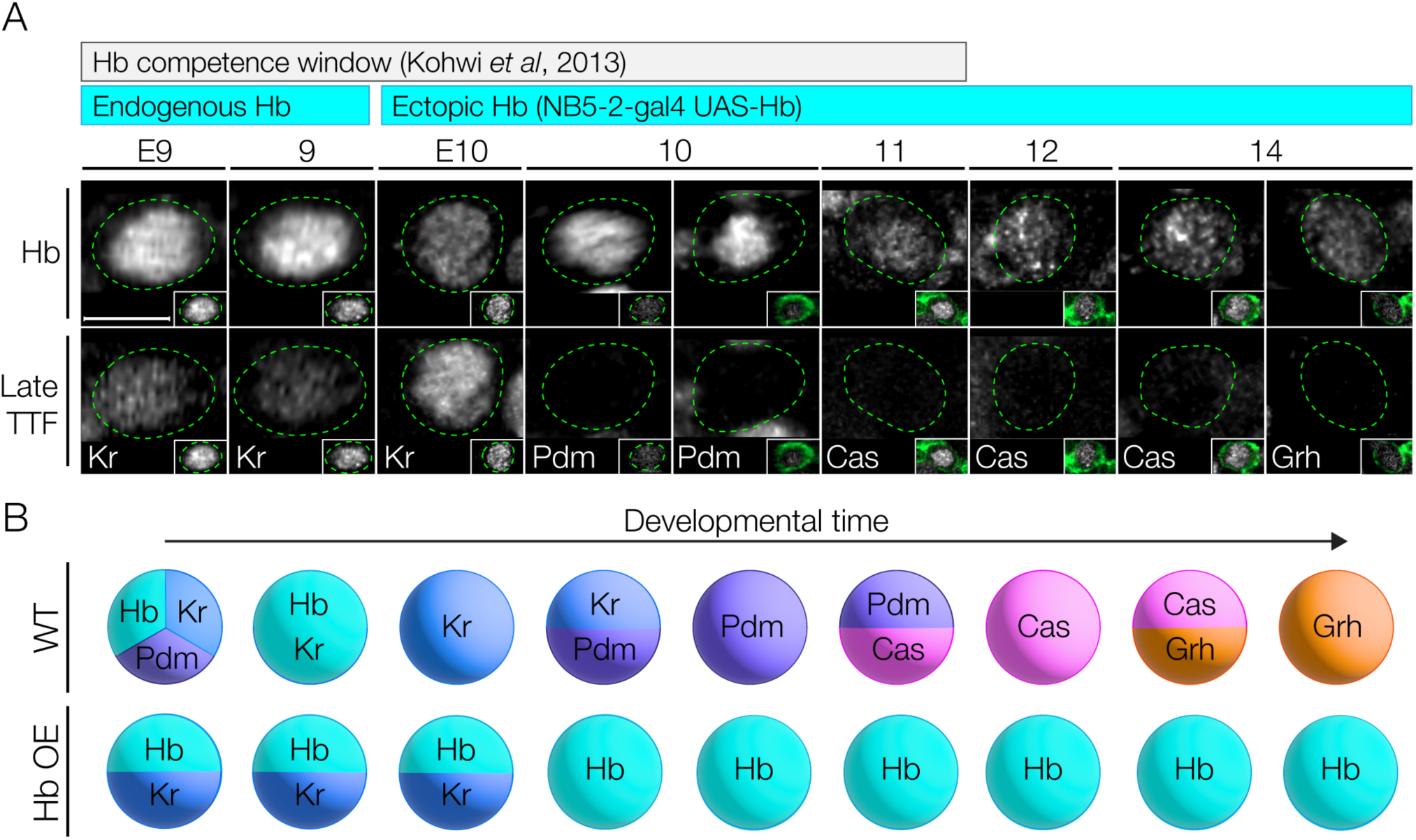
Prolonged Hunchback expression in NB5-2 delays expression of late TTFs. (A) NB5-2>Hb progeny Hb (top panels) and late TTF (bottom panels) expression at stages 9-14. Bars show developmental length of known Hb competence window in other NBs (gray) and endogenous and ectopic NB5-2 Hb expression (cyan). NB5-2 identification was achieved using *dpn* and *en* in stages 9 and early stage 10 embryos (see SuppFig 1); a combination of GFP (green) and *dpn* was used following stage 10 (insets). Scale bar: 5 µm (B) Schematic of wildtype NB5-2 progressing through the TTF cascade: Hb (cyan)>Kr (blue)>Pdm (purple)>Cas (magenta)>Grh (orange), compared to NB5-2>Hb (HbOE).

### Hunchback specifies early-born interneuron molecular identity

Next, we wanted to determine if loss of later-born TTFs due to prolonged Hb expression extended to NB5-2 progeny. We quantified the number of NB5-2 neurons expressing each TTF following Hb misexpression. As expected, we observed a significant increase in the number of Hb+ neurons in the NB5-2 lineage (Figure 5E-F, quantified in Q). We also found an increase in Hb+/Kr+, and Hb/+Pdm+ co-expressing neurons (Supp. Figure 2A-B, quantified in D-G), consistent with co-expression observed in wildtype NB5-2 neurons (Figure 2A). We note that while NB Pdm expression is repressed by Hb misexpression, we see an increase in Hb+/Pdm+ NB5-2 progeny; the discrepancy in Pdm expression from NB to progeny is unclear. Conversely, we observed a striking decrease in Cas+ and Grh+ neurons, of which NB5-2 Hb+ neurons have no overlapping expression (Figure 5K-L, quantified in U; Supp. Figure 2C, quantified in H). We conclude that NB5-2>Hb is an effective tool for prolonging Hb expression throughout the NB5-2 lineage, and that prolonged Hb expression is highly effective at repressing NB expression of all later TTFs.

**Figure 5.**
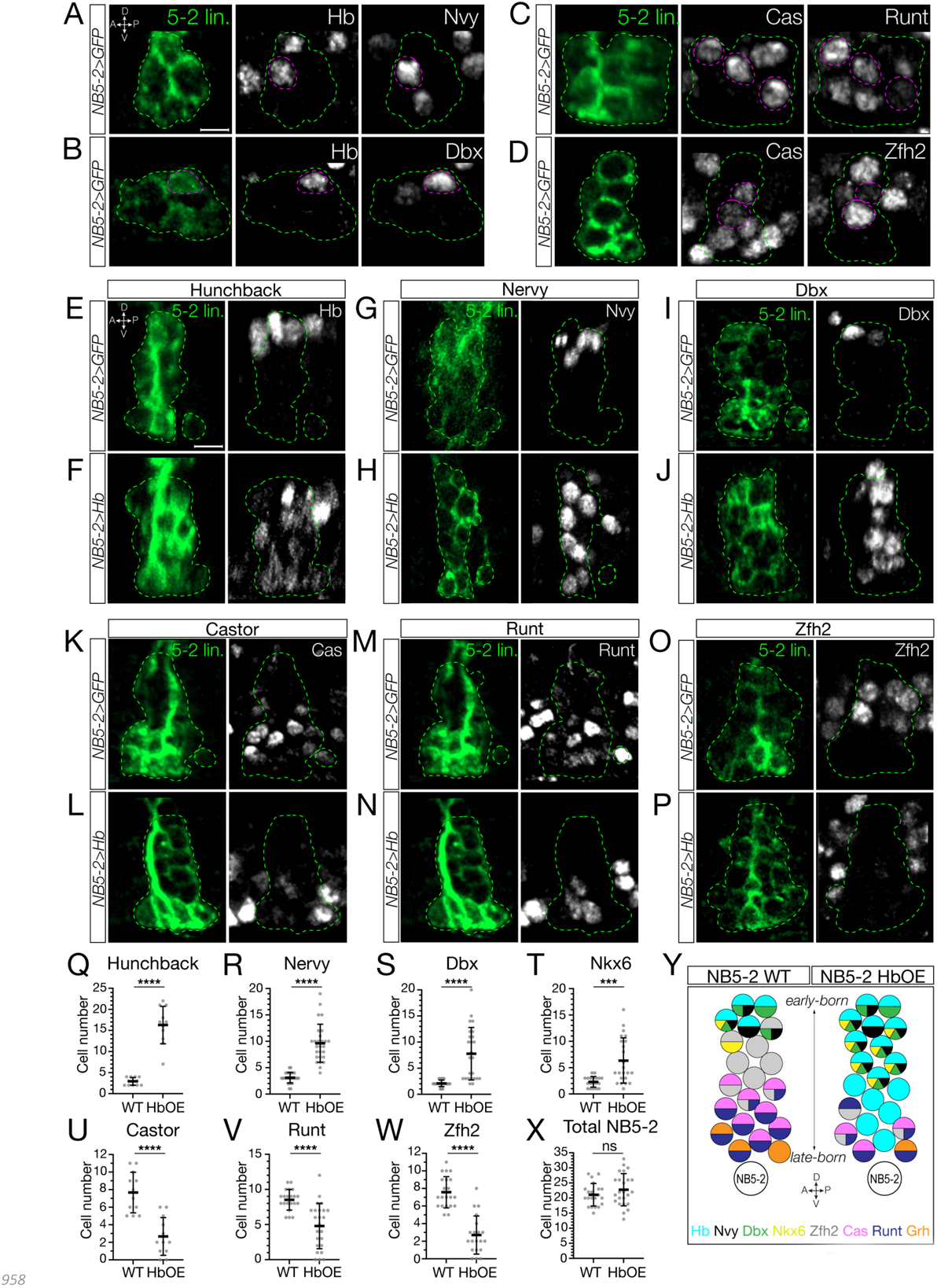
Prolonged Hunchback expression in NB5-2 generates ectopic early-born interneurons at the expense of late-born neurons. (A) Wildtype NB5-2>GFP (WT; green) progeny co-expressing Hb and the early-born TF, Nervy, at stage 17 (dotted magenta). Anterior left, lateral view. Scale bar: 4 µm. (B) WT NB5-2>GFP progeny co-expressing Hb and the early-born TF, Dbx. (C) WT NB5-2>GFP progeny co-expressing Cas and the late-born TF, Runt. (D) WT NB5-2>GFP progeny co-expressing Cas and the late-born TF, Zfh2. (E) WT NB5-2 progeny expressing Hb at stage 16/17. Anterior left, lateral view. Scale bar: 4 µm. (F) NB5-2>Hb progeny expressing Hb. (G) WT NB5-2>GFP progeny expressing Nervy (Nvy). (H) NB5-2>Hb progeny expressing Nervy. (I) WT NB5-2>GFP progeny expressing Dbx. (J) NB5-2>Hb progeny expressing Dbx. (K) WT NB5-2>GFP progeny expressing Cas. (L) NB5-2>Hb progeny expressing Cas. (M) WT NB5-2>GFP progeny expressing Runt. (N) NB5-2>Hb progeny expressing Runt. (O) WT NB5-2>GFP progeny expressing Zfh2. (P) NB5-2>Hb progeny expressing Zfh2. (Q) Quantification of WT NB5-2>GFP (avg=2.75, n=12 hemisegments, 3 animals) and NB5-2>Hb (HbOE; avg=16.27, n=11 hemisegments, 3 animals) neurons expressing Hb at stage 16/17 (*p-value*<0.0001). (R) Quantification of WT NB5-2 (avg=3.11, n=28 hemisegments, 7 animals) and HbOE (avg=9.63, n=27 hemisegments, 7 animals) neurons expressing Nervy (*p-value*<0.0001). (S) Quantification of WT NB5-2 (avg=2.09, n=22 hemisegments, 6 animals) and HbOE (avg=7.76, n=15 hemisegments, 4 animals) neurons expressing Dbx (*p-value*<0.0001). (T) Quantification of WT NB5-2 (avg=2.31, n=19 hemisegments, 5 animals) and HbOE (avg=6.35, n=23 hemisegments, 6 animals) neurons expressing Nkx6 (*p-value*=0.0003). (U) Quantification of WT NB5-2 and HbOE neurons expressing Cas (WT avg=7.67, HbOE avg=2.67, n=12 hemisegments, 3 animals, *p-value*<0.0001). (V) Quantification of WT NB5-2 (avg=8.52, n=21 hemisegments, 6 animals) and HbOE NB5-2 (avg=4.79, n=24 hemisegments, 6 animals) neurons expressing Runt (*p-value*<0.0001). (W) Quantification of WT NB5-2 (avg=7.57, n=21 hemisegments, 6 animals) and HbOE NB5-2 (avg=2.7, n=20 hemisegments, 6 animals) neurons expressing Zfh2 (*p-value*<0.0001). (X) Quantification of total WT (avg=21.10, n=21 hemisegments, 6 animals) and HbOE (avg=22.75, n=24 hemisegments, 6 animals) cell number in NB5-2 lineage (*p-value*=0.24; n.s). (Y) Schematic summary of NB5-2 WT and HbOE cells, expressing the TTFs Hb (cyan), Cas (magenta), and Grh (orange); the early-born TFs, Nervy (Nvy; black), Dbx (green), and Nkx6 (yellow); the late-born TFs, Zfh2 (grey) and Runt (purple).

To determine the role of Hb in specifying interneuron <u>molecular identity</u> downstream of TTF expression, we needed to identify molecular markers for early-born Hb+ and late-born Hb-negative interneurons in the NB5-2 lineage. We screened a collection of transcription factor antibodies, and identified Nervy, Dbx, and Nkx6 as early-born TFs expressed in Hb+ neurons but not in Cas+ neurons (Figure 5A,B and Supp. Figure 3A). Conversely, we identified Runt and Zfh2 as late-born TFs expressed in Cas+ neurons, but not in Hb+ neurons (Figure 5C,D). Next, we prolonged Hb expression in the NB5-2 lineage and assayed for changes in TF expression. We found that Hb misexpression resulted in an expansion of early-born Hb+, Nervy+, Dbx+, and Nkx6+ neurons that spanned the deep-superficial axis of the NB5-2 lineage (Figure 5E-J and Supp. Figure 3B,C; quantified in Q-T). Furthermore, prolonged Hb expression resulted in a striking loss of late-born Cas+, Runt+, and Zfh2+ neurons (Figure 5K-P; quantified in U-W). Importantly, there was no significant difference in the total number of NB5-2 neurons per hemisegments (Figure 5X), showing that late-born neurons are transformed into neurons with an early-born molecular identity, rather than undergoing cell death. We conclude that Hb is sufficient to specify early-born interneuron molecular identity at the expense of late-born neurons, and that temporal identity is more important than birth-order in specifying early-born interneuron molecular identity.

### Hunchback+ early-born interneurons have a distinct morphology compared to later-born interneurons

To determine the role of Hb in specifying NB5-2 early-born interneuron <u>axon/dendrite morphology</u>, we needed to identify the individual neuron morphology of both early-born Hb+ and late-born Hb-negative interneurons. We showed above that NB5-2 makes five Hb+ neurons (Figure 2). Here we design a genetic method for identifying the morphology of each Hb+ neuron in the NB5-2 lineage. We used intersectional genetics to stochastically label individual neurons that were both Hb+ and derived from NB5-2. We call this intersectional approach “Hb+ filtered NB5-2 neurons” (Figure 6A). Using this method, we identified five Hb+ neurons in the NB5-2 lineage: MN12 (Figure 6B), and four interneurons that we name “Idun1-Idun4” (Figure 6C-F). Each Idun neuron has a similar but unique morphology. Based on deep-superficial position, we propose that the first NB5-2 derived GMC (GMC-1) makes MN12/Idun1 siblings, GMC-2 makes Idun2/Idun3 siblings, and GMC-3 makes Idun4 and a sibling that may undergo apoptosis, as we have never detected a sixth Hb+ neuron in the lineage. Subsequently we focus on the four Hb+ Idun interneurons.

**Figure 6.**
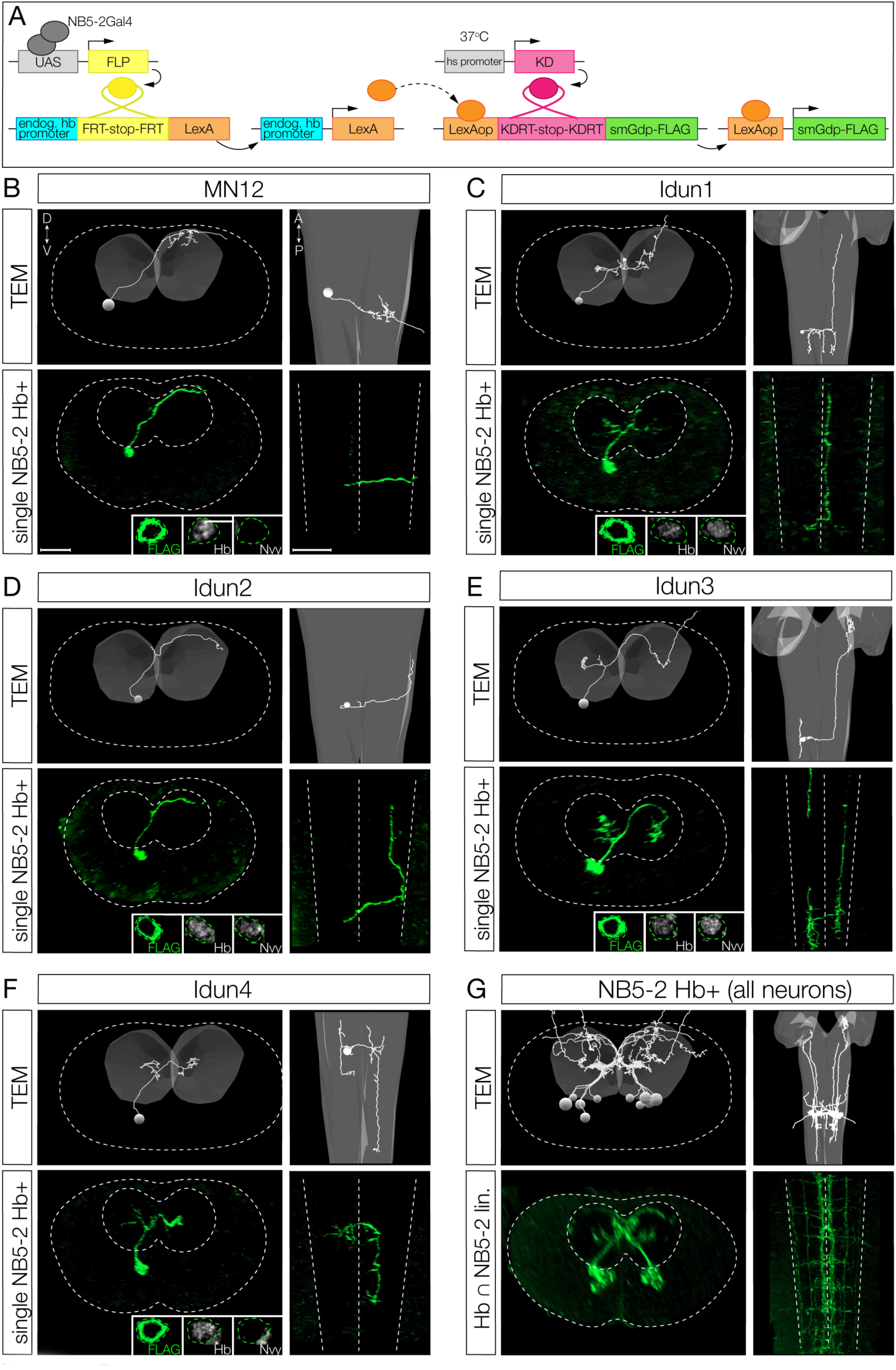
Hunchback+ early-born interneurons have a distinct morphology compared to later-born neurons. (A) Intersectional genetics used to stochastically label NB5-2 Hb+ neurons. NB5-2 Gal4 (grey) drives expression of Flipase (FLP, yellow) to promote excision of a stop codon inserted downstream of the endogenous Hb promoter (cyan) and upstream of a LexA transgene (orange). Heat-stocking embryos drives expression of KD recombinase (magenta) to promote excision of a stop codon downstream of a LexAop site. In combination of LexA expression, this allows for expression of spaghetti monster with a FLAG epitope tag (green) exclusively in NB5-2 Hb expressing neurons. (B) Motor neuron 12 (MN12) morphology traced using transmission electron microscopy (TEM; top panels) and labeled using Hb+ filtered NB5-2 intersectional genetics (single NB5-2 Hb+; bottom panels). MN12 cell body labeled with FLAG (green; left inset) shows Hb (middle inset) expression but lacks Nervy (Nvy; right inset) expression. Dorsal up, posterior view (left panels); Anterior up, ventral view (right panels). Scale bar: 10 µm (C) Idun1 interneuron morphology traced using TEM (top panels) and labeled single NB5-2 Hb+ neuron (bottom panels). Idun1 shows Hb (middle inset) and Nvy (right inset) expression. (D) Idun2 interneuron morphology traced using TEM (top panels) and labeled single NB5-2 Hb+ neuron (bottom panels). Idun2 shows Hb (middle inset) and Nvy (right inset) expression. (E) Idun3 interneuron morphology traced using TEM (top panels) and labeled single NB5-2 Hb+ neuron (bottom panels). Idun3 shows Hb (middle inset) and Nvy (right inset) expression. (F) Idun4 interneuron morphology traced using TEM (top panels) and labeled single NB5-2 Hb+ neuron (bottom panels). Idun4 shows Hb (middle inset) expression but lacks Nvy (right inset) expression. (G) TEM reconstruction (top panels) and Hb+ ^∩^ NB5-2 lin. labeling (bottom panels) of all Hb+ NB5-2 neurons.

We previously used the open-source transmission electron microscopy (TEM) volume of the first instar larval CNS [29] to reconstruct all of the neurons within the NB5-2 lineage in both segment A1L and A1R [11], but it was unknown which neurons in the whole lineage reconstruction were Hb+. To identify the morphology of the Hb+ neurons in the lineage, we used the intersectional genetics method described above to label “Hb+ filtered NB5-2 neurons” with a membrane-bound reporter (Figure 6A). We were able to identify all four Hb+ Idun interneurons as well as MN12 based on a clear match in neuron morphology (Figure 6B-F). All Hb+ Idun interneurons are unique, but there are a few shared features. (1) All Idun interneurons have a cell body position in a deep layer close to the neuropil. (2) All Idun interneurons have an ascending or descending intersegmental projection. (3) With the exception of Idun4, all Idun interneurons have a contralateral projection that crosses the midline with a diagonal trajectory, from ventral-medial to dorso-lateral, clearly visible in a posterior view.

We found that the formation of a diagonal contralateral projection was unique to the early-born neurons; TEM reconstruction of all neurons in the lineage only found this trajectory in Idun1, Idun2, Idun3 and MN12. These shared morphological features can be observed when all five Hb+ neurons in the lineage are labeled together by light or electron microscopy (Figure 6G). In conclusion, we have identified four Hb+ interneurons in the NB5-2 lineage and matched them to the same four interneurons within the TEM reconstruction. Importantly, these early-born interneuron morphologies are clearly different from late-born interneuron morphology (Supp. Figure 4). In addition to identifying morphological differences in early-born and late-born interneurons, mapping the Idun interneurons in the TEM volume allows us to quantify presynapse and postsynapse position for each Idun neuron - a prerequisite for characterization of Idun presynapse neuropil targeting (see below).

### Hunchback specifies early-born interneuron morphology

We showed above that Hb determines early-born interneuron molecular identity based on early and late-born TF expression. Here we ask whether Hb determines early-born interneuron <u>axon/dendrite</u> <u>morphology</u>. We wanted to distinguish between two possible mechanisms, as we did for interneuron molecular identity. First, *birth-order* may be critical for determining neuronal morphology, perhaps each neuron born at a different time encounters a changing environment which directs the appropriate projection. Second, *TTF expression* may determine neuronal morphology, perhaps via TTF-regulated guidance cues. To determine the relative importance of birth-order and TTF expression, we misexpressed Hb in the NB5-2 lineage and asked whether late-born neurons persisted in generating late-born neuronal morphology (birth-order model) or whether they acquired early-born neuronal morphology (TTF model). The latter was found to be important in the motor neurons of the NB7-1 lineage [14,15]; here we ask whether this mechanism is extended to the interneurons, where axons and dendrites remain within the synaptically dense neuropil of the CNS.

We focus on Idun1, Idun2, and Idun3, as these interneurons have a unique diagonal crossing projection not observed in later-born neurons, with the addition of Idun1 possessing the most medial ascending projection and Idun2 possessing the most lateral ascending projection among NB5-2 progeny (Figure 7). NB5-2>GFP labels three types of contralateral projections: dorsal, ventral, and diagonal, that are best seen in a posterior view (Figure 7A), plus ascending/descending projections that are best seen in a dorsal view (Figure 7A’). In contrast to controls, Hb misexpression shows a striking loss of the dorsal and ventral contralateral projections and an increase in the diagonal projections (Figure 7B; posterior view) as well as a loss of ascending/descending projections (Figure 7B’). These phenotypes are due to a change in late-born neuron morphology, and not the death of late-born neurons, as there are the same number of neurons in control and experimental NB5-2 lineages (Figure 5X).

**Figure 7.**
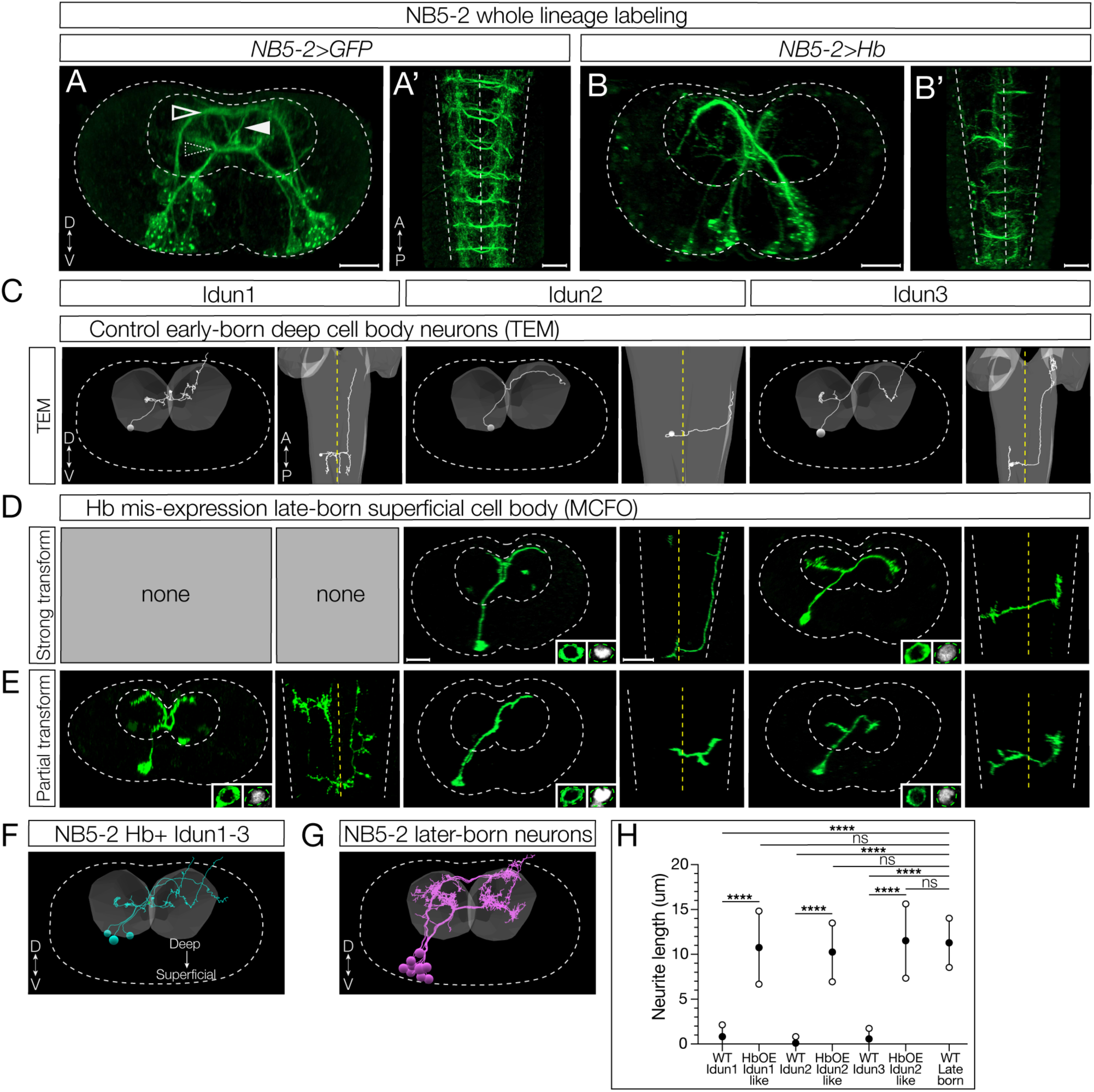
Hunchback determines early-born neuron morphology. (A) Whole lineage labeling of wildtype (WT) NB5-2>GFP (green) axon/dendrite morphology displaying dorsal (outlined arrow), ventral (dotted arrow), and diagonal (filled arrow) neurite projections within the A1/A2 VNC neuropil. Dorsal up, posterior view; Anterior up, ventral view (A’). Scale bar: 10 µm. (B) Whole lineage labeling of NB5-2>Hb morphology within the A1/A2 VNC neuropil. Dorsal up, posterior view; Anterior up, ventral view (B’). (C) TEM reconstruction of WT Idun1 (left two panels), Idun2 (middle two panels), and Idun3 (right two panels) morphology and cell body position relative to the neuropil (grey). Dorsal up, posterior view (left panels of each neuron); Anterior up, ventral view (right panels of each neuron). VNC cortex border (white dotted line); VNC midline (yellow dotted line). (D) NB5-2>Hb late-born neurons, labeled using MultiColor FlipOut (MCFO; green), displaying strong Idun1 (left two panels), Idun2 (middle two panels), and Idun3 (right two panels) morphology with Hb (right inset) expression in the cell body (left inset). Dorsal up, posterior view (left panels of each neuron); Anterior up, ventral view (right panels of each neuron). Scale bar: 10 µm. (E) NB5-2>Hb late-born neurons displaying partial Idun1 (left two panels), Idun2 (middle two panels), and Idun3 (right two panels) morphology and Hb (right inset) expression in the cell body (left inset). (F) TEM reconstruction of WT NB5-2 Hb+ neuron morphology from a single A1 hemisegment. (G) TEM reconstruction of WT NB5-2 later-born neuron morphology from a single A1 hemisegment. (H) Quantification of WT Idun1-3, late-born transformed Idun1-3 NB5-2>Hb (HbOE Idun1-like/Idun2-like/Idun3-like), and WT late-born neurite length (WT Idun1 avg=0.83, n=8; WT Idun2 avg=0.11, n=7; WT Idun3 avg=0.58, n=5; HbOE Idun1-like avg=10.76, n=9; HbOE Idun2-like avg=10.25, n=10; HbOE Idun3-like avg=11.52, n=8; WT late-born avg=11.29, n=19; one-way ANOVA *p-values*: WT Idun1 vs HbOE Idun1-like *p-value* <0.0001, WT Idun2 vs HbOE Idun2-like *p-value* <0.0001, WT Idun3 vs HbOE Idun3-like *p-value* <0.0001, WT Idun1 vs. WT late-born *p-value*<0.0001, WT Idun2 vs. WT late-born *p-value*<0.0001, WT Idun3 vs. WT late-born *p-value*<0.0001, HbOE Idun1-like vs. WT late-born *p-value*=0.99 (n.s), HbOE Idun2-like vs. WT late-born *p-value*=0.97 (n.s), HbOE Idun3-like vs. WT late-born *p-value*>0.99 (n.s)).

To prove that Hb misexpression reprograms late-born interneurons to generate an axon/dendrite morphology characteristic of early-born neurons, we labeled single neurons in the NB5-2 lineage using multicolored flip out (MCFO) [30] in a wild type or Hb misexpression background. Following Hb misexpression, we identified late-born interneurons based on their superficial cell body position (see Figure 3) [11], and confirmed that they were Hb+ despite their superficial position (Figure 7D-E). We found that these late-born neurons expressing Hb had an early-born axon/dendrite morphology, including a diagonal contralateral projection and ascending/descending intersegmental projections similar to the TEM reconstruction of Idun1-3 neurons (Figure 7C-E); we call these “strongly” transformed neurons. In addition, we also observed “partially” transformed late-born neurons, characterized by the expected diagonal commissural projection but lacking ascending or descending projections (Figure 7E). Why there are two phenotypic classes is not clear (see Discussion).

To gain additional evidence for the Hb-mediated transformation of late-born neurons into neurons with morphological features characteristic of early-born neurons, we measured soma-neuropil distance as a proxy for birth-order (see Figure 3) [11]. As expected, wild type early-born Idun1, Idun2, and Idun3 neurons had cell bodies close to the neuropil, with very short soma-neuropil distance (Figure 7F, quantified in H). Also as expected, wild type Hb-negative late-born neurons had superficial locations and a large soma-neuropil distance (Figure 7G, quantified in H). Importantly, Hb misexpression resulted in neurons with a diagonal commissural projection, characteristic of early-born neurons, yet showing a large soma-neuropil distance (Figure 7H). This shows that late-born, superficially located neurons can be efficiently transformed into neurons with early-born morphology following prolonged Hb expression. We conclude that Hb is sufficient to induce early-born interneuron morphology. This is consistent with the “TTF model” for specifying interneuron morphology.

### Hunchback determines interneuron presynapse targeting within the neuropil

We next wanted to determine if Hb played a role in early-born interneuron <u>presynaptic targeting</u>, which is particularly interesting due to the dense packing of synapses in the neuropil. We first sought to determine the presynapse localization of Idun1-3 within the neuropil, a critical first step in establishing proper connectivity to downstream early-born partners. We used the TEM dataset to determine the neuropil volume occupied by all NB5-2 neuronal presynapses (Figure 8A) as well as specifically for Idun1, Idun2, and Idun3 (Figure 8B). The neuropil position of presynapses differed for each neuron, which we label as <u>N</u>europil-1 (N1), N2 and N3 for Idun1-3, respectively. We then mapped the coordinates for the three volumes to the neuropil and used Hb filtered genetics (Figure 8C) to express the presynaptic marker Bruchpilot (Brp) specifically in the Idun1-3 neurons. Importantly, in controls the Brp puncta targeted the same three volumes in both light and electron microscopy, which can be seen for both Hb+ neurons (Figure 8D) and all NB5-2 progeny neurons (Figure 8E). Misexpression of Hb results in several phenotypes. (1) There are more Brp puncta targeting N1 and N2 (compare Figure 8E and F; quantified in Figure 8G), consistent with late-born neurons being transformed to early-born Idun1 and Idun2 neurons, and targeting to their characteristic Idun1/2 neuropil volumes. (2) There is a decrease in Brp puncta in the N3 volume. (3) There are Brp puncta abnormally positioned between the N2 and N3 volumes (Figure 8F; yellow arrows). Existing tools do not allow us to further understand the second and third phenotypes (see Discussion), but the first phenotype provides clear support for a model in which the early TTF Hb can reprogram late-born neurons, driving them to target their presynapses to neuropil volumes normally targeted by presynapses of early-born neurons.

**Figure 8.**
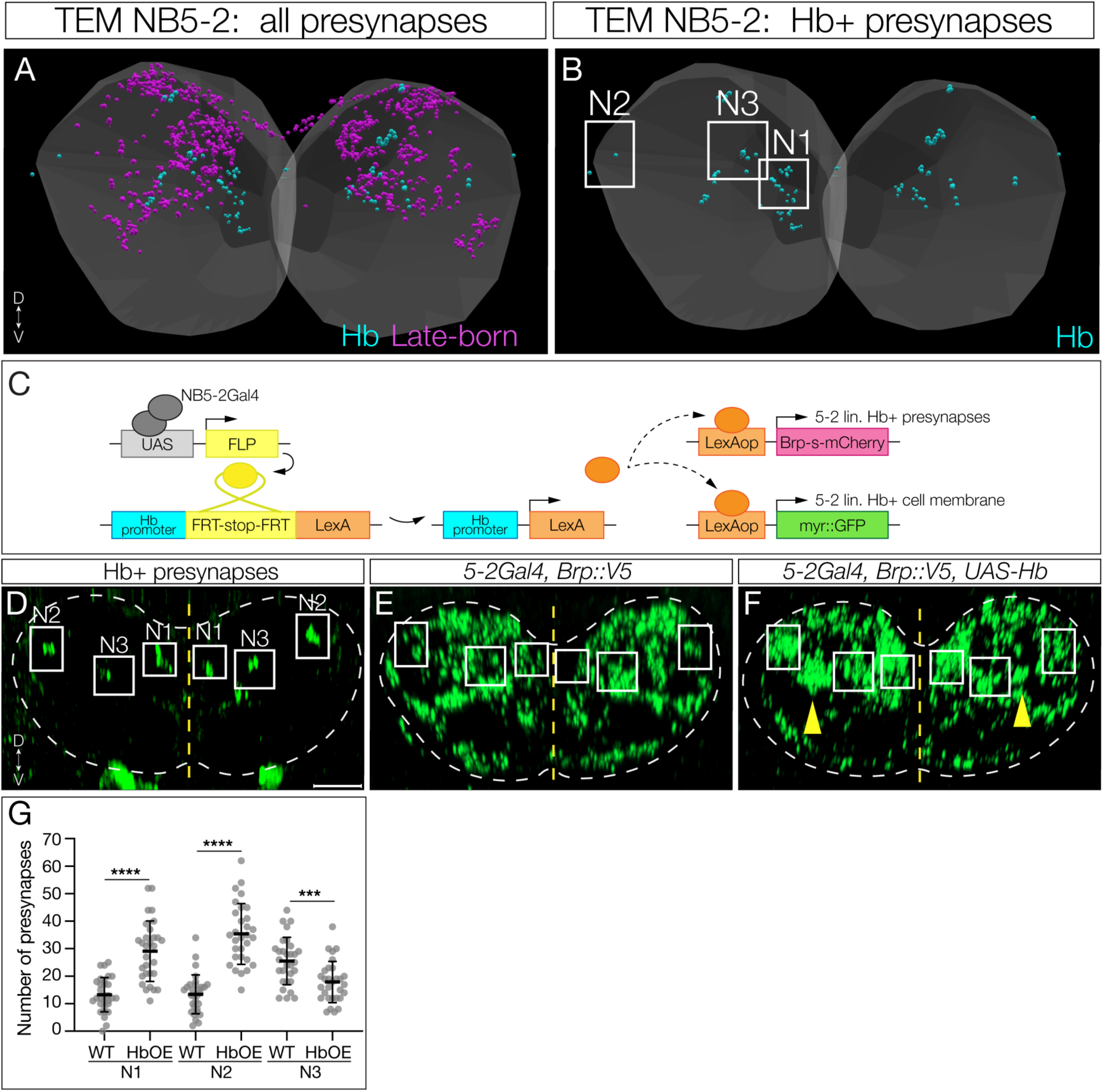
Hunchback determines interneuron presynapse localization. (A) TEM reconstruction of NB5-2 Hb+ (cyan) and late-born (magenta) presynapses in the neuropil (grey). Dorsal up, posterior view. (B) TEM reconstruction of NB5-2 Hb+ Idun1 (<u>N</u>europil position 1, N1), Idun2 (<u>N</u>europil position 2, N2) and Idun3 (<u>N</u>europil position 3, N3) presynapse positions in the left and right A1 hemisegments. (C) Intersectional genetics approach to drive expression of Brp exclusively in NB5-2 Hb expressing neurons. NB5-2 Gal4 (grey) drives expression of Flipase (FLP, yellow) to promote excision of a stop codon inserted downstream of the endogenous Hb promoter (cyan) and upstream of a LexA transgene (orange), allowing for LexA expression exclusively in NB5-2 Hb expressing neurons. LexA then drives expression of a truncated Brp conjugated to mCherry (Brp-s-mCherry, magenta) and GFP (myr::GFP, green). (D) NB5-2 Hb+ presynapses labeled by Brp expression (green) using intersectional genetics. Idun1-3 positions are outlined (white) in the neuropil (dotted outline) according to TEM reconstruction. Dorsal up, posterior view. Scale bar: 5 µm. (E) WT NB5-2 presynapses labeled by endogenous Brp expression conjugated to a V5 epitope tag (green). Idun1-3 positions are outlined. (F) NB5-2>Hb presynapses with Idun1-3 positions outlined and a possible Idun3 mistargeting location (yellow arrow; see Discussion). (G) Quantification of WT NB5-2 and NB5-2>Hb (HbOE) larval presynapses in neuropil positions N1 (WT avg=13.27, HbOE avg=29.10, n=30, *p-value*<0.0001), N2 (WT avg=13.43, HbOE avg=35.37, n=30, *p-value*<0.0001), and N3 (WT avg=25.50, HbOE avg=17.9, n=30, *p-value=*0.0006).

### Hunchback misexpression phenocopies loss of a proprioceptive circuit

*Drosophila* larvae have a well-characterized proprioceptive circuit: proprioceptive sensory neurons > Jaam1-3 > Eve+ interneurons > Saaghi1-3 > motor neurons [23,31]. Ablation of the Eve+ interneurons in the proprioceptive circuit results in decreased larval crawling speed and increased C-shaped body bends [23]. Importantly, both Jaams and Saaghis are late-born neurons in the NB5-2 lineage [11]. This raises the question: if Hb misexpression transforms late-born Jaam and Saaghi neurons into an early-born identity, will they maintain or lose their connectivity to the proprioceptive circuit neurons? If Hb misexpression can drive late-born neurons into a circuit with endogenous early-born neurons, we would expect a loss of proprioceptive behavior. Thus, we misexpressed Hb throughout the NB5-2 lineage and measured these two behaviors. Qualitatively, experimental larvae showed a strong uncoordinated crawling pattern (compare SuppMovie 1 and SuppMovie 2). Quantitatively, we observed significantly decreased forward locomotor velocity and increased C-shaped body bends, compared to controls (Figure 9A,B). We conclude that transformation of late-born NB5-2 interneurons to an early-born identity disrupts late-born proprioceptive circuit assembly or function.

**Figure 9.**
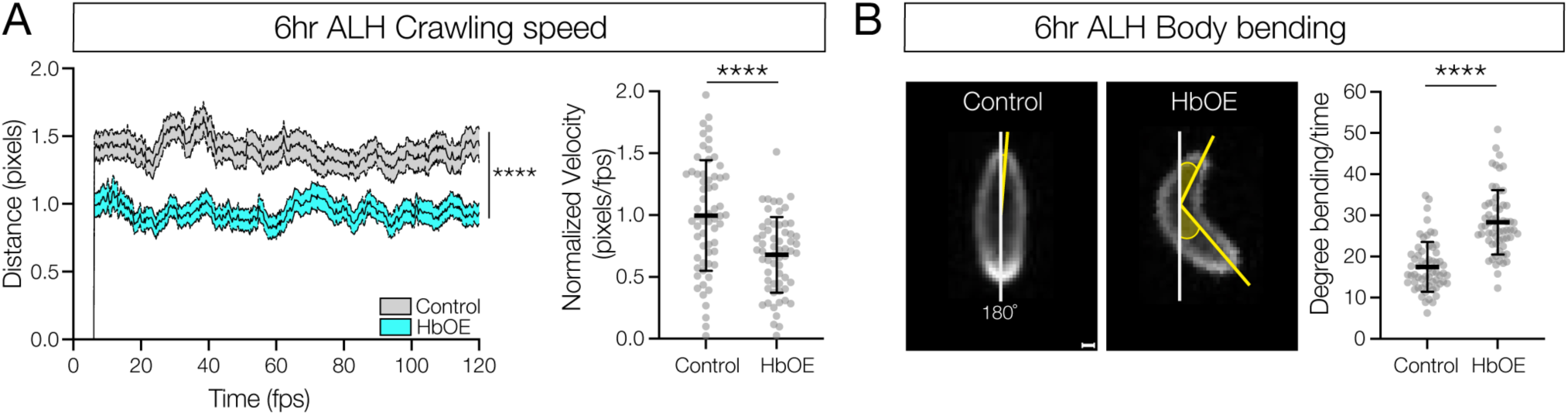
Prolonged Hunchback expression disrupts late-born proprioceptive circuit connectivity. (A) Crawling speed of newly hatched WT NB5-2>GFP (Control; grey, avg AUC=159, n=61)) and NB5-2>Hb (HbOE; cyan, avg AUC=106.9, n=62) larvae (0-4hrs ALH) (*p-value*<0.0001); Normalized crawling speed of WT (control, avg=0.99) and HbOE (avg=0.68) larvae (*p-value*<0.0001). (B) WT NB5-2 (control, avg=17.47, n=61) and HbOE (avg=28.31, n=62) body bend posture determined by the angle of degree (yellow shaded) from 180° (white line); Quantification of body-bend degree from 180° (*p-value*<0.0001). Scale bar: 40 µm.

## Discussion

During neurogenesis, intrinsic mechanisms play a vital role in determining neuron identity. The role of Hb in specifying motor neuron identity has been well studied in NB3-1 and NB7-1 [9,13,17,32–34]. Our study provides a comprehensive analysis of Hb in specifying NB5-2 interneuron identity. We found that the TTF cascade is expressed sequentially in NB5-2 (Hb > Kr > Pdm > Cas > Grh) and is transiently maintained in the post-mitotic progeny with single or multiple gene overlap combinations. This expression pattern is consistent with the previously characterized NB3-1 and NB7-1 and is further support of a conserved, intrinsic TTF mechanism among VNC NBs [2]. To determine the role of Hb in specifying interneuron identity we prolonged Hb expression in NB5-2 and analyzed three aspects of neuron identity: <u>molecular identity</u>, <u>axon/dendrite morphology</u>, and <u>presynapse targeting</u> (required for proper connectivity), and then assessed behavioral output due to changes in neuron identity.

We found that prolonged Hb expression prevented expression of late-born TTFs and increased the number of interneuron progeny with an early-born molecular identity (Nervy+ Dbx+ Nkx6+) at the expense of late-born progeny. In NB3-1, prolonged Hb expression increases the number of RP1/RP4 motor neurons with an early-born Zfh2-/Cut-molecular identity [12,16]. Similarly, NB7-1 Hb misexpression increases motor neuron progeny with an early-born U1/U2 molecular identity (Eve+/Zfh2-) [14,15]. In NB5-6 and NB7-4, both of which generate interneuron progeny, Hb binds NB specific loci, suggesting differential expression of downstream genes [35]. Previous work has also shown that the late TTF Cas window of expression can be subdivided by “sub-temporal” genes to specify four distinct Apterous+ (Ap+) interneurons in the NB5-6T lineage. Ap+ neurons are uniquely specified through activation of the transcription factor Collier/Knot (Col) and a series of feedforward loops [36,37]. The combinatorial expression of Kr/Pdm has also been shown to be crucial for the Nkx6+ molecular identity of the VO MN, derived from NB7-1 [38]. Thus, the combinatorial expression of multiple TTFs may be important in promoting individual neuronal molecular identities. In the NB5-2 lineage, Hb induced expression of the early-born TFs (Nervy, Dbx, and Nkx6) suggests that Hb may promote specification of distinct interneurons through differential expression of unique TF combinations.

To understand if Hb promotes NB5-2 interneuron morphology, we first identified the unique morphology of the NB5-2 Hb+ interneurons, Idun1-4, and then found that late-born neurons displayed a strikingly similar early-born diagonal neurite morphology following Hb misexpression. We propose three explanations for these phenotypic classes: (1) partially transformed late-born neurons do not experience the same developmental environment as early-born neurons and thus lack the environmental signals to extend ascending and descending projections. (2) Late-born neurons inherently have less time to form long projecting neurites. (3) Neurons may require a separate mechanism to form long projecting neurites, not present in some late-born neurons.

Prolonged Hb expression in NB7-1 results in a late-born to early-born transformation of U1/U2 early-born dendrite morphology [14,16,16]. The expression of extrinsic signaling ligand/receptors, such as Sema/Plexin or Slit/Robo, allows a neuron to respond to a given attractant or repulsive signaling gradient within the neuropil [22]; differential expression of one or more receptor in early- or late-born neurons may determine differences in presynaptic or postsynaptic neuropil targeting. A transcriptomic study of *Drosophila* olfactory projection neurons (PNs) revealed that transcriptomes of PN subtypes shows the highest amount of TF and cell-surface molecule diversity during circuit assembly [39]. In future studies, identifying and functionally testing downstream Hb targets such as guidance ligands/receptors will be an important next step in understanding how connectivity within the *Drosophila* CNS is established.

To understand if Hb promotes functional connectivity, we would ideally like to optogenetically simulate Idun1-3 pre-synaptic partners (or Idun1-3 directly) following Hb misexpression. A TTX-insensitive increase in GCaMP activity in putative downstream partners would functionally validate monosynaptic connectivity. Unfortunately, TEM connectivity data revealed that many Idun2 and Idun3 pre- and post-synaptic partners are unknown neurons, meaning we do not currently possess genetic access to these neurons. Idun1 shows pre- and post-synaptic connections to the well characterized Moonwalking Descending Neuron (MDN) [40–43]. The connectivity between MDN and Idun1 is the most ideal connection among the NB5-2 Hb+ neurons because we have genetic access to MDN, and Idun1 expresses GABA, suggesting it functions as an inhibitory neuron (Supp. Figure 5); however, there are two additional late-born NB5-2 neurons that also form synapses with MDN, making an excitation response between endogenous and transformed late-born neurons impossible to distinguish.

Although genetic limitations prevented us from testing whether Hb transformed late-born neurons acquire functional connectivity matching that of early-born neurons, we performed two experiments that both support a Hb-induced switch in late-born neuron connectivity. First, we asked if Hb misexpression in late-born neurons induces them to form presynapses in neuropil domains normally targeted by endogenous early-born neurons. We found a significant increase in presynapse number in Idun1 and Idun2 subregions following Hb misexpression, consistent with the transformed neurons acquiring the same connectivity as the endogenous early-born neurons. However, we also observed a significant decrease in presynapses in the N3 volume and an abnormally positioned cluster of presynapses between N2 and N3 (Figure 8F; yellow arrows). These results may be consistent with Idun3 being transformed to Idun1 or Idun2; alternatively, the Idun3 presynapse region contains some late-born presynapses, and thus it is possible that these neurons are transformed into Idun1/2 identity. In the future, direct genetic assess to Idun1-3 will allow us to explore these possibilities.

Second, we asked if Hb misexpression in late-born neurons disrupted their normal connectivity, leading to slow locomotion and increased C-shaped body bends, hallmarks of defective proprioceptive behavior [23]. Indeed, Hb misexpression resulted in the same behavioral defects as ablation of proprioceptive circuit neurons, consistent with reprograming late-born neuron connectivity to neurons of the proprioceptive circuit to a new connectivity that is unable to generate normal proprioceptive behavior. These findings are consistent with Hb misexpression abolishing normal late-born neuron connectivity. Note that the role of TTFs in establishing connectivity has been previously shown for embryonic motor neurons [14–16] and there are strong correlations between neuronal birth-order and wiring specificity in *Drosophila* olfactory projection neurons and mushroom body Kenyon cells [44].

In *Caenorhabditis elegans*, the *hb* homolog, *hbl-1*, specifies early-born seam cell fate [45]. In addition, combinatorial expression of homeodomain proteins specify specific neuron types [46]. Our data shows that Hb is sufficient to promote the expression of two homeodomain proteins, Nkx6 and Dbx, in early-born NB5-2 progeny, and repress homeodomain Zfh2 expression. Hb repression of Zfh2 is also observed in early-born NB3-1 and NB7-1 neurons [12,14–16]. Taken together, this may place Hb as a hierarchical transcription factor that promotes/represses the expression of downstream homeodomain proteins, which then generate unique, lineage specific TF combinations to specify distinct aspects of individual neuron identity.

In both flies and mammals, neuron settling position is correlated with neuronal birth-order. For example, early-born Hb+ neurons are located deep in the VNC cortex and late-born Cas+ are located superficially. In mammals, cortical projection neurons in different layers project to distinct regions of the CNS [47,48]. Progeny generated from apical radial glia (aRG) settle in a similar birth-order dependent manner, early-born neurons located in deep cortical layers while later-born neurons migrate to settle more superficially [1]. The Hb mammalian ortholog, Ikaros (Ikzf1), is expressed early in aRGs and is necessary and sufficient to generate early-born progeny [6,7]. Prolonged Ikzf1 expression in aRGs increases the number of deep layer early-born cells, identified by early-born TFs (Ctip2+Tbr1+Foxp2+) [7]. Similarly, Ikzf1 is expressed in early retinal progenitor cells (RPC) and promotes the production of early-born neuron identities derived from RPCs. Misexpression of Ikzf1in RPCs increases the number of early-born horizontal cells and amacrine cells at the expense of late-born bipolar neurons [6]. In addition, the Cas mammalian ortholog, Casz1, is expressed in later cell divisions of RPCs and promotes the production of late-born rod cells [49,50].

Our data shows that Hb expression in NB5-2 is sufficient to specify early-born interneuron identity at the molecular, morphological, and presynapse targeting levels. The correlations in *Drosophila* NBs and mammalian aRGs and RPCs suggests there is a conserved mechanism in which intrinsic TTF expression in progenitor cells is, in part, an initial requirement to diversify individual progenitor lineages. This leads to many open questions: how does Hb promote lineage-specific early-born neuron identity? What is the function of TFs downstream of Hb? Is the role of Hb in specifying interneuron identity conserved across multiple species? Does Ikzf1 promote/repress the expression of specific TFs important for distinct aspects of early-born cortical or retinal cell identity? These will be exciting avenues to explore in future studies.

## Materials and Methods

### Fly Stocks

Male and female *Drosophila melanogaster* were used. The chromosomes and insertion sites of transgenes (if known) are shown next to genotypes.

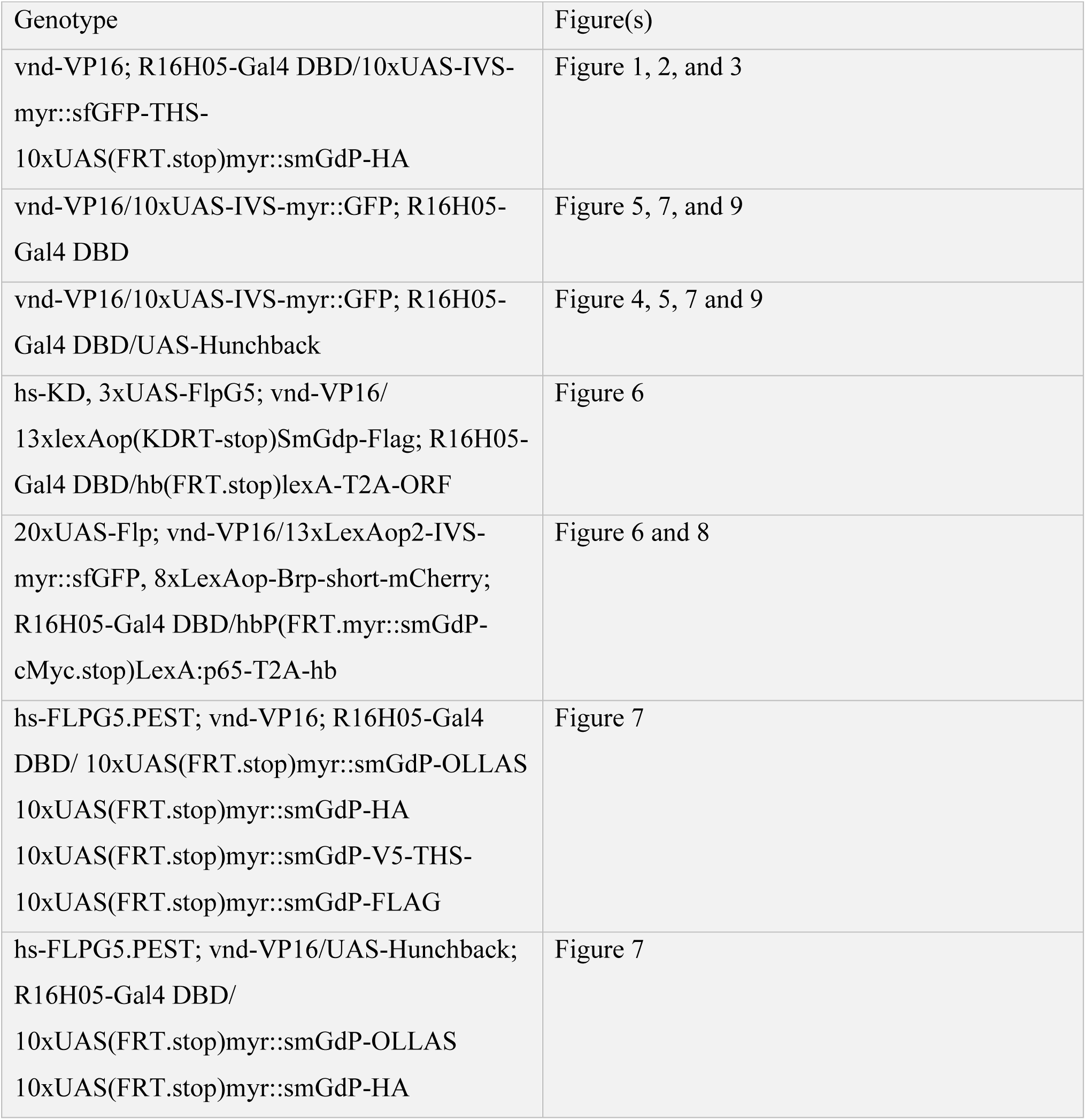
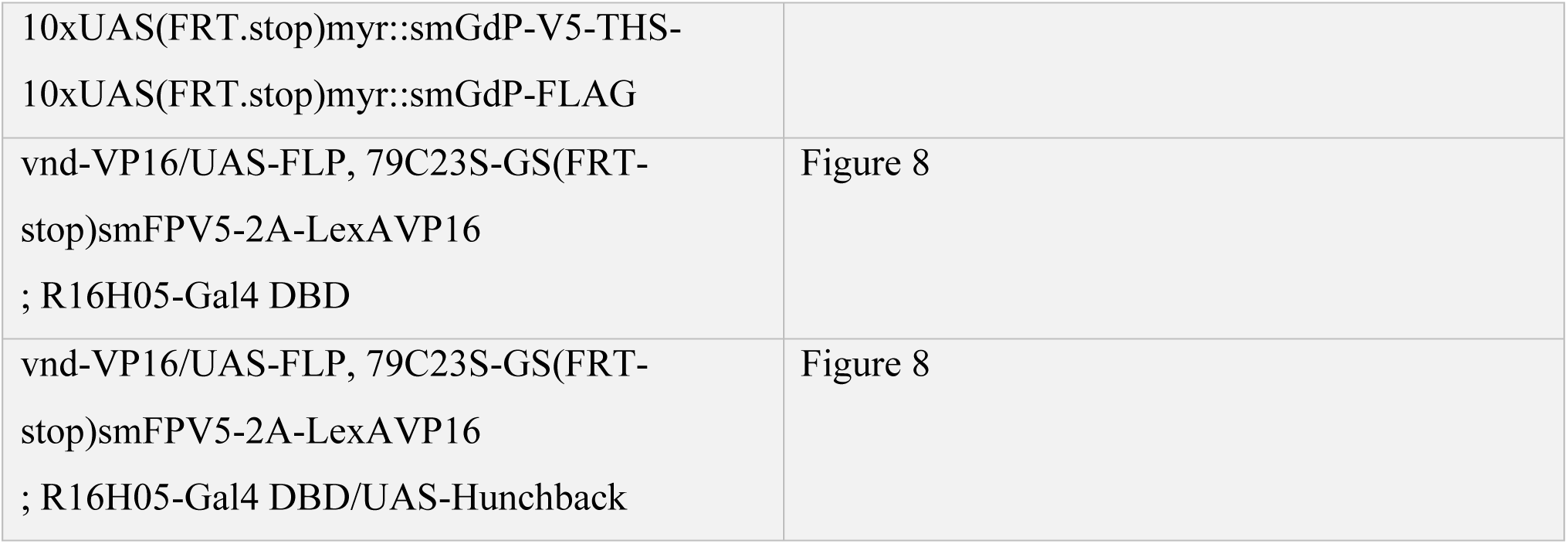

Gal4 lines, mutants and reporters used were:

vnd-VP16 (2R, VK1, 59D3) [11]

R16H05-Gal4 DBD (3L, P2, 68A4) (this work)

10XUAS-IVS-myr::sfGFP-THS-10xUAS(FRT.stop)myr::smGdP-HA (RRID:BDSC_62127)

10xUAS-IVS-myr::GFP (BDSC_32198)

UAS-Hunchback (III) [9]

hs-KD (BDSC_56167)

3xUAS-FLPG5 (BDSC_55808)

13xlexAop(KDRT-stop)SmGdp-Flag (BDSC_62111)

hbP(FRT.myr::smGdP-cMyc.stop)LexA:p65-T2A-hb [11]

hs-FLPG5.PEST.Opt (RRID:BDSC_77140)

10xUAS(FRT.stop)myr::smGdP-OLLAS 10xUAS(FRT.stop)myr::smGdP-HA

10xUAS(FRT.stop)myr::smGdP-V5-THS-10xUAS(FRT.stop)myr::smGdP-FLAG

(RRID:BDSC_64086)

20xUAS-FLP (BDSC_55807)

13xLexAop2-IVS-myr::sfGFP (this work)

8xLexAop-Brp-short-mCherry [51]

UAS-FLP (BDSC_4540)

79C23S-GS(FRT-stop)smFPV5-2A-LexAVP16 [52]

### Neuron naming

We named the Hb+ interneurons in the NB5-2 lineage Idun1-4 after Iðunn, the Norse god of youth.

### Immunostaining and imaging

Primary antibodies were: chicken anti-GFP (1:1000, RRID:AB_2307313, Aves Labs, Davis, CA), mouse anti-Hunchback (1:100, Abcam, F18-1G10.2), rabbit anti-Hunchback #5-27 (1:400) [12], guinea pig anti-Kr (1:500, Doe lab), rat anti-Pdm2 (1:100, abcam, ab201325, Cambridge, MA), rabbit anti-Castor (1:1000, Doe lab), rat anti-Grainy head (1:400, a gift from Stefan Thor, University of Queensland), rat anti-Deadpan (1:20, abcam, ab195173), rabbit anti-Worniu (1:1000; abcam, ab196362), rabbit anti-Nervy (1:300, gift from Richard Mann, Columbia University), rat anti-Nkx6 (1:500, gift from J. Skeath, Washington University, St. Louis), guinea pig anti-Runt (1:1000, gift from C. Desplan, New York University), rat anti-Zfh2 (1:250, Doe lab), rat anti-FLAG (1:400, Novus NBP1-06712), rat anti-HA (1:1000, MilliporeSigma, 11867423001, St. Louis, MO), chicken anti-V5 (1:800, Bethyl Laboratories, Inc. A190-118A, Centennial, CO), rabbit anti-mCherry (1:500, Novus, NBP2-25157), mouse anti-Eve (5ug/mL, DSHB 2B8), rat anti-N-cadherin (0.168ug/mL, DSHB DN-Ex #8), and fluorophores-conjugated secondary antibodies were from Jackson ImmunoResearch (West Grove, PA) and were used at 1:300 for embryos and 1:400 for larval brains.

Embryos were fixed in 4% paraformaldehyde or formaldehyde for 20 min and stained as previously described [11]. Larval brains were dissected in PBS, fixed in 4% paraformaldehyde, and then stained by following protocols as described [53]. The samples were DPX mounted.

Images were captured with a Zeiss LSM 900 confocal microscope with a *z*-resolution of 0.22 μm. Due to the complex 3-dimensional pattern of each marker assayed, we could not show NB5-2 progeny marker expression in a maximum intensity projection, because irrelevant neurons in the z-axis obscured the neurons of interest; thus, NB5-2 progeny were montaged from their unique z-axis position while preserving their X-Y position. Images were processed using Imaris (Bitplane, Zurich, Switzerland) to generate three dimensional reconstructions and level adjustments. Any level adjustment was applied to the entire image. Figures were assembled in Adobe Illustrator (Adobe, San Jose, CA).

### Presynapse analysis

Neuropil volume was labeled using N-Cadherin antibody staining. To define neuropil borders we made a surface object using Imaris 10.0.1 software. We then defined a centroid in each hemisegment by measuring the length, height, and depth of a single abdominal hemisegment, focusing our analysis on segments 1 and 2 (A1 and A2) (Supp. Figure 6). For Hb+ neuropil volumes N1-N3, presynapses distribute along the anterior-to-posterior axis, spanning the depth of a single hemisegment. To control for segmental depth, we used the transcription factor marker Even-skipped (Eve) to locate the U1 motor neurons and defined segmental borders according to their location (i.e. the U1 motor neuron position in thoracic segment 3 defined the A1 anterior border; the U1 position in A1 define the posterior border). To isolate individual presynapses, we used the Imaris 10.0.1 spots tool and found the average center of each Hb+ position (N1 n=10 animals, 20 hemisegments; N2 n=10 animals, 20 hemisegments; N3 n=4 animals; 9 hemisegments). To find the distance of each center from the centroid, we measured and averaged the distances along the medial-lateral and dorsal-ventral axes from the centroid. A volume for each position was then created using Imaris 10.0.1 custom surface tool, spanning the depth of the hemisegment. The size of each volume was determined by the average center distance plus/minus 2 standard deviations. Presynapses that fell within each volume where then quantified by hand.

### Behavior

We recorded behavior in newly hatched larvae (4-6hr). Behavior arenas were made with 1.2% agar, 2 mm thick. Arenas were placed on a Functional Independence Measure (FIM) table for recording. Larvae were transferred to the agar arena and given 2 min to acclimate before recording. Behavior was recoded for 2 min (125 fps) at 24°C and 60% humidity using a Basler acA2040-25gm camera with a Computer TEC-V7X 1.1” 7X Macro Zoom Telecentric lens and Pylon Viewer software. Recordings were edited in Fiji to eliminate arena borders, adjust the brightness-to-contrast ratio to increase larvae visibility (Min:15-19, Max:57-64), and saved at 25fps. Larval locomotion speed and body bends were calculated using FIMTrack 2.0 software.

### Statistics

Statistical significance is denoted by asterisks: ****p<0.0001; ***p<0.001; **p<0.01; *p<0.05; n.s., not significant. The following statistical tests were performed: Welch’s t-test (normal distribution, non-parametric) (two-tailed p value) (Figures 5Q-X, 8G, 9A-B) and one-way ANOVA (Figures 7H). All analyses were performed using Prism8 (GraphPad). The results are stated as mean ± s.d., unless otherwise noted.

## Acknowledgements

We thank fellow lab member Noah Dillon for constructive comments on the manuscript. We also thank Tory Herman (Oregon) for comments on the manuscript. We thank Sen-Lin Lai and Chundi Xu for reagents and the construction of fly genetics. We thank Stefan Thor, Richard Mann, Jim Skeath, and Claude Desplan for antibody reagents. We thank Kristen Lee for training and feedback on behavior experiments. Antibodies obtained from the Developmental Studies Hybridoma Bank, created by the NICHD of the NIH and maintained at the University of Iowa, Department of Biology, Iowa City, IA were used in this study. Stocks obtained from the Bloomington Drosophila Stock Center (NIH P40OD018537) and Vienna Drosophila Resource Center were used in this study. Funding was provided by HHMI and NIH HD27056 (CQD) and T32 HD07348 (HQP).

---

**Supplemental Figure 1.**
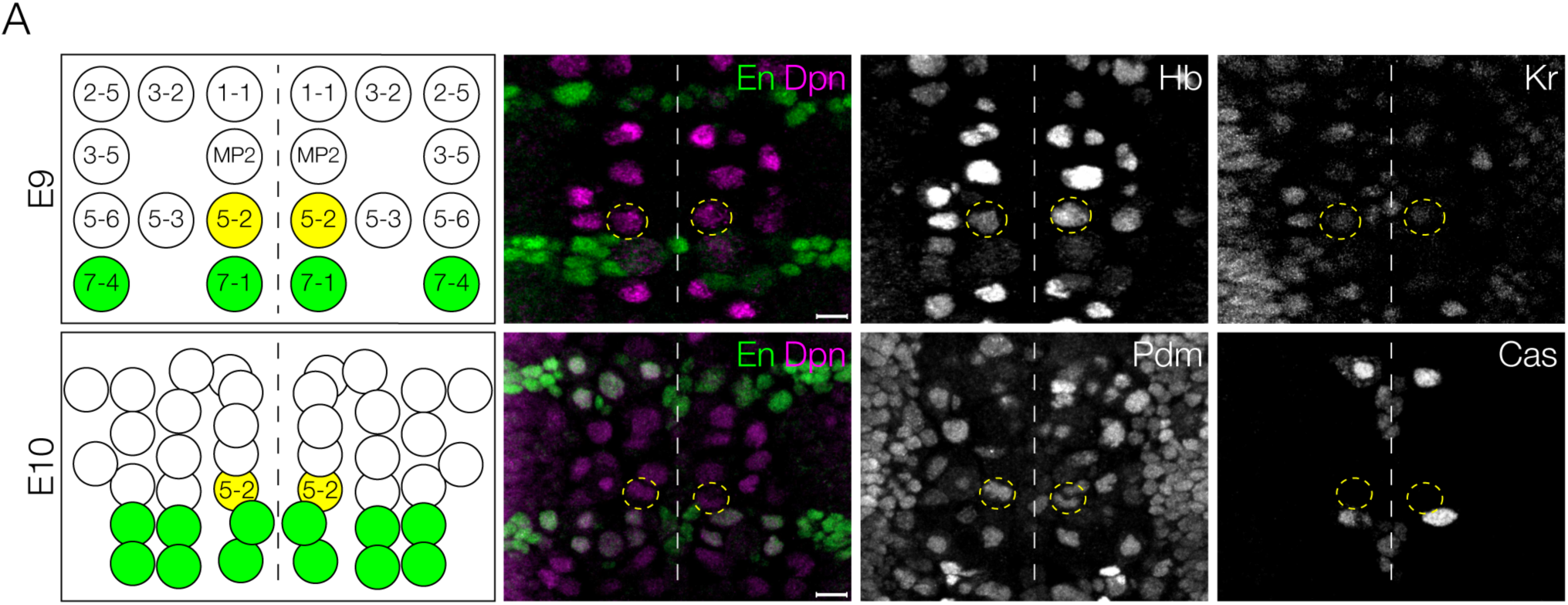
NB5-2 can be identified by its stereotyped location in early embryos. NB5-2 (yellow in schematic) was identified using the NB marker *dpn* (magenta) and the row 6/7 expressing gene, *engrailed* (*en*; green). NB5-2 was identified as the most medial dpn+/en-negative NB, anteriorly adjacent to the engrailed domain. NB5-2 shows Hb/Kr expression in early stage 9 embryos (top panels) and Pdm expression by early stage10 (bottom panels). Scale bar: 5 µm.

**Supplemental Figure 2.**
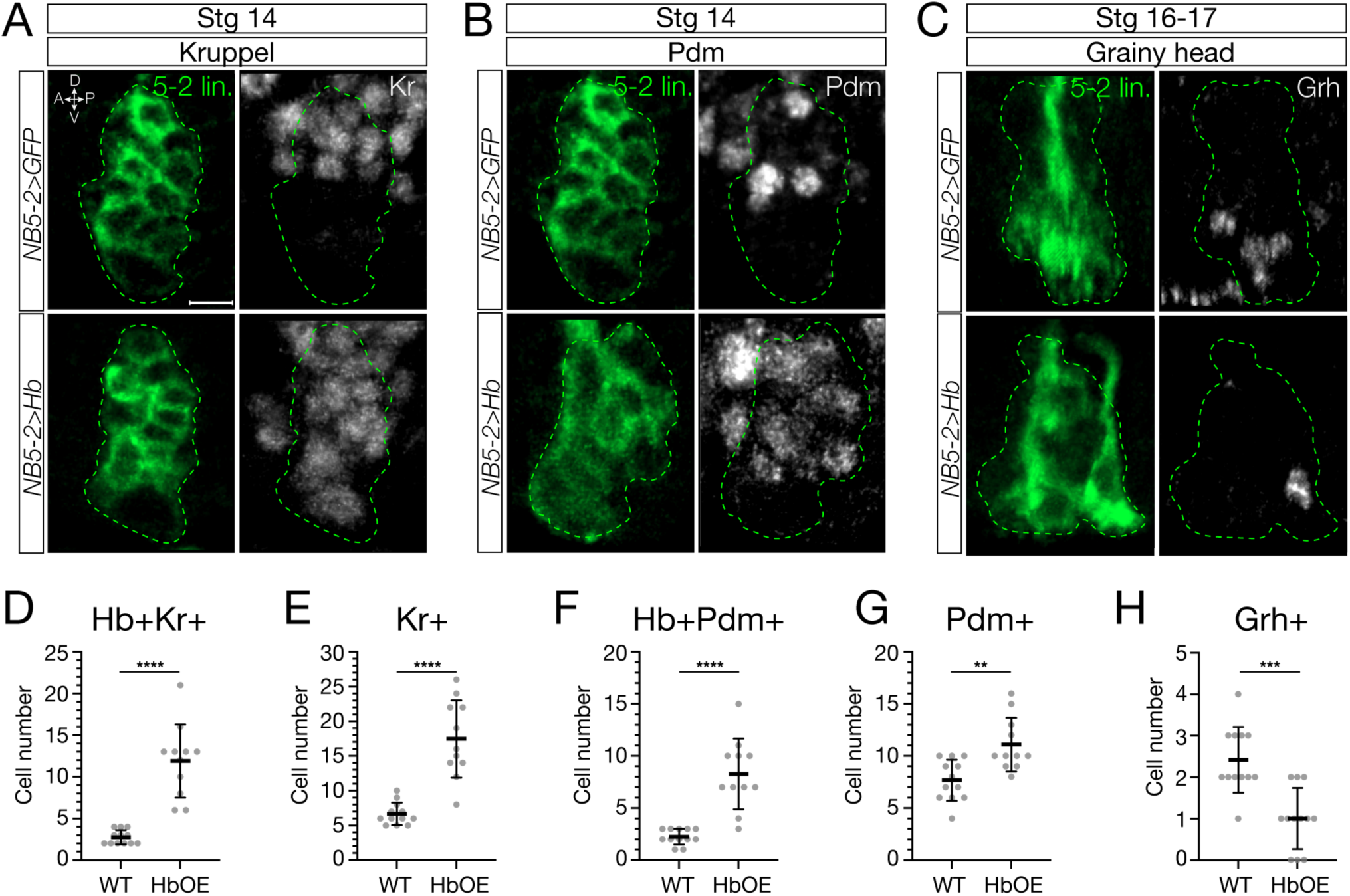
Prolonged Hunchback expression in NB5-2 generates fewer progeny expressing late TTFs. (A) Wildtype NB5-2>GFP (green; top panels) and NB5-2>Hb (bottom panels) progeny expressing Kr at stage 14. Anterior left, lateral view. Scale bar: 4 µm. (B) Wildtype NB5-2>GFP (top panels) and NB5-2>Hb (bottom panels) progeny expressing Pdm at stage 14. (C) Wildtype NB5-2>GFP (top panels) and NB5-2>Hb (bottom panels) progeny expressing Grh at stage 16-17. (D) Quantification of wildtype NB5-2>GFP (WT; avg=2.75, n=12 hemisegments, 3 animals) and NB5-2>Hb (HbOE; avg=11.91, n=11 hemisegments, 4 animals) neurons expressing Hb and Kr (*p-value*<0.0001). (E) Quantification of all wildtype NB5-2>GFP (avg=6.67, n=12 hemisegments, 3 animals) and NB5-2>Hb (avg=17.45, n=11 hemisegments, 4 animals) neurons expressing Kr (*p-value*<0.0001). (F) Quantification of wildtype NB5-2>GFP (avg=2.25, n=12 hemisegments, 3 animals) and NB5-2>Hb (avg=8.27, n=11 hemisegments, 3 animals) neurons expressing Hb and Pdm (*p-value*<0.0001). (G) Quantification of all wildtype NB5-2>GFP (avg=7.67, n=12 hemisegments, 3 animals) and NB5-2>Hb (avg=11.09, n=11 hemisegments, 3 animals) neurons expressing Pdm (*p-value*=0.0017). (H) Quantification of all wildtype NB5-2>GFP (avg=2.42, n=12 hemisegments, 3 animals) and NB5-2>Hb (avg=1.00, n=12 hemisegments, 3 animals) neurons expressing Grh (*p-value*=0.0002).

**Supplemental Figure 3.**
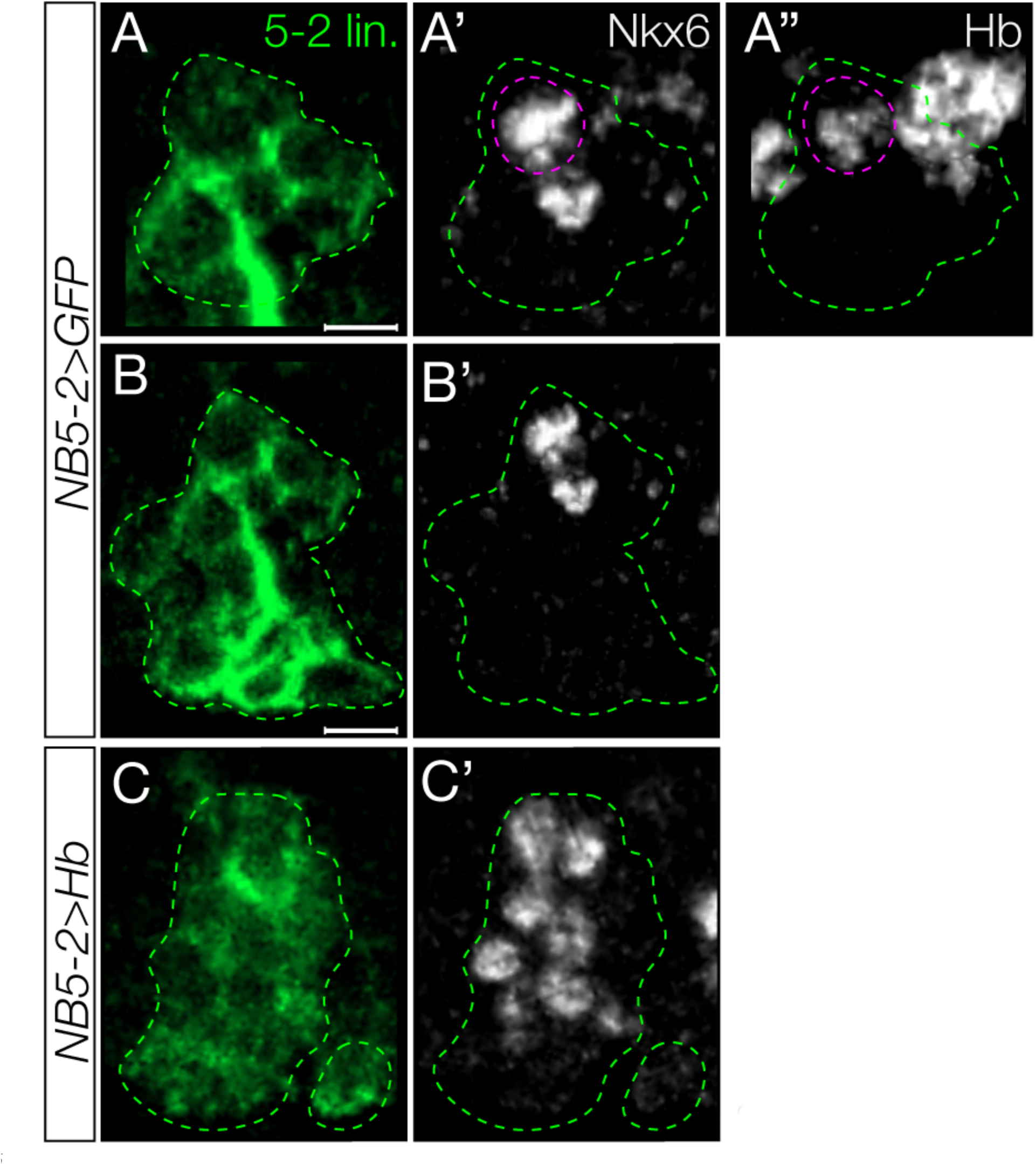
Prolonged Hunchback expression increases the number of NB5-2 progeny that express Nkx6. (A) Wildtype NB5-2>GFP (WT; green) progeny co-expressing Hb and the early-born TF, Nkx6, at stage 17 (dotted magenta). Anterior left, lateral view. Scale bar: 4 µm. (B) WT NB5-2>GFP progeny expressing Nkx6 (B’), at stage 17. Scale bar: 4 µm. (C) NB5-2>Hb progeny expressing Nkx6 (C’).

**Supplemental Figure 4.**
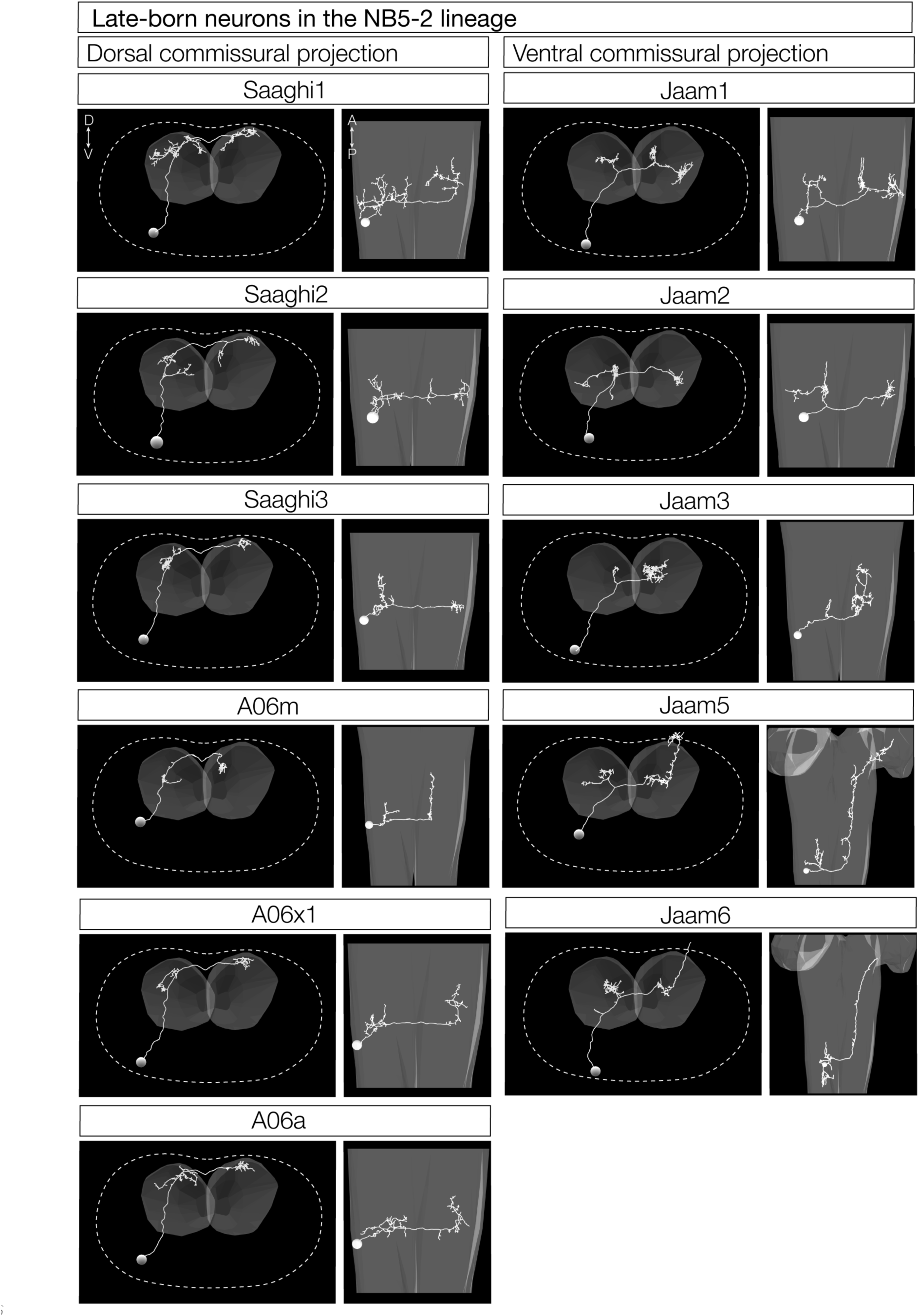
NB5-2 late-born neurons do not possess a diagonal projecting morphology. TEM reconstruction of wildtype NB5-2 late-born neurons organized by dorsal commissural projections (left column) and ventral commissural projections (right column). Dorsal up, posterior view (left panel); Anterior up, ventral view (right panel).

**Supplemental Figure 5.**
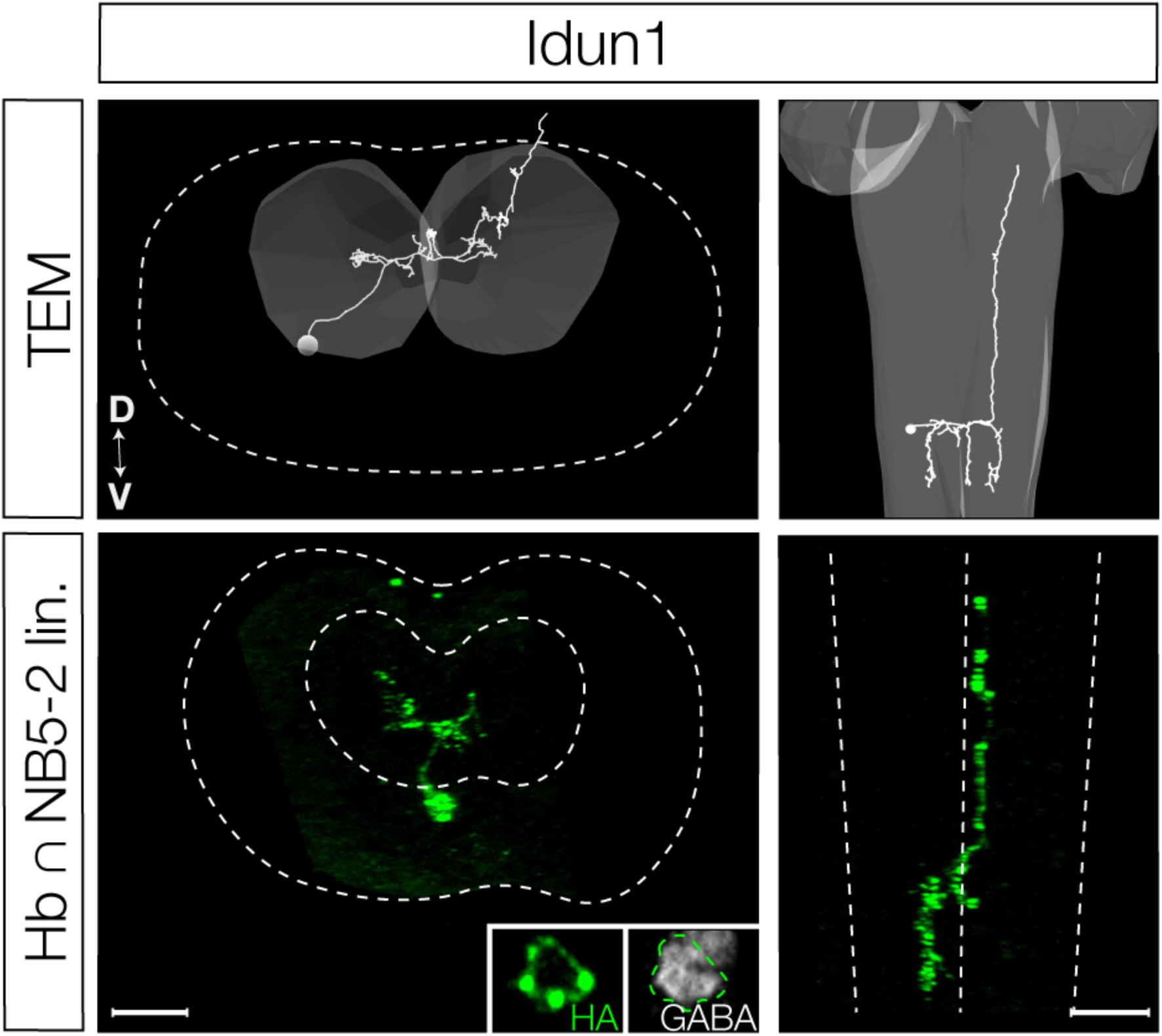
Idun1 expresses GABA neurotransmitter. TEM reconstruction of Idun1 (upper panels) and single labeled Hb+ NB5-2 neuron genetically labeled with membrane-bound epitope tag, HA (green; bottom panels), with GABA expression (inset). Dorsal up, posterior view (left panels); Anterior up, ventral view (right panels). Scale bar: 10 µm

**Supplemental Figure 6.**
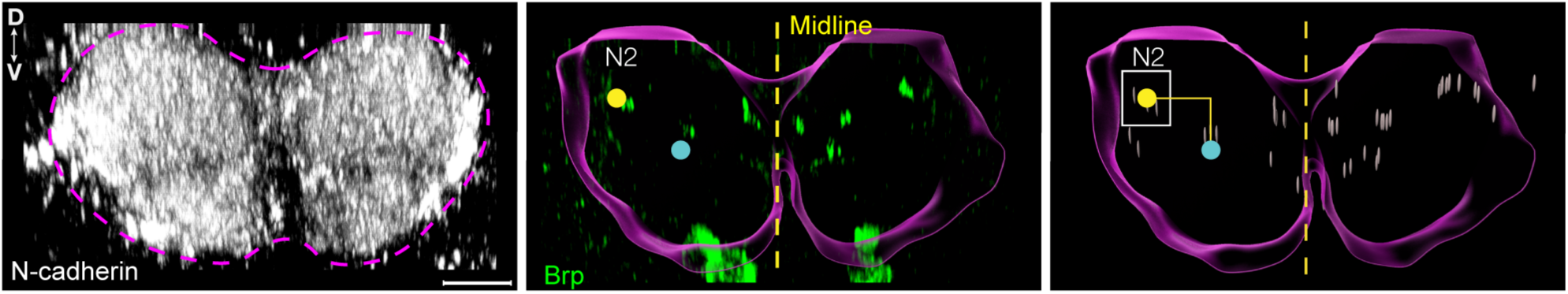
Approach to identify NB5-2 Hb+ presynapse neuropil subregions. Neuropil labeled with N-cadherin (white) to define the neuropil border (magenta dotted line; left panel). The neuropil border (magenta) was defined using Imaris 10.0.1 surface tool to find the centroid (cyan dot) of a hemisegment and the center of Idun1-3 presynapse position labeled with Brp staining (green; middle panel). Shown is the Idun2 presynapse neuropil position (N2; yellow dot). The average N2 coordinate location was then found by measuring the dorsal and lateral distance from the centroid (yellow solid line; right panel). The size of the presynapse volume (white box) was determined by the average distance from the centroid +/- 2 standard deviations. Presynapses were simplified using the Imaris 10.0.1 spots tool (grey dots) for quantification. Dorsal up, posterior view. Scale bar: 5 µm.

